# Spatial inhibition of RhoA by RhoGAP15B promotes protrusive activity during collective migration

**DOI:** 10.64898/2025.12.30.694567

**Authors:** Vitor Yang, Cristian Marchant, Bing Liu, Mariana Osswald, Jesus Lopez-Gay, Yohanns Bellaiche, Xiaobo Wang, Eurico Morais-de-Sá

## Abstract

The Rho family GTPases RhoA, Rac1 and Cdc42 are well established regulators of collective migration by driving the formation of cellular protrusions and by regulating actomyosin contraction and adhesion. However, how their activation and inhibition are spatially and temporally coordinated remains unclear. Using GFP knock-in lines, we systematically characterized the localization patterns of all *Drosophila* RhoGEFs (activators) and RhoGAPs (inhibitors) in border cells, an *in vivo* model of collective migration. We have further combined RNAi screening with GFP-based validation of depletion efficiency to assess the functional significance of those RhoGEF/GAPs expressed in border cells. This identified RhoGAP15B as a localized inhibitor of RhoA activity at the border cell cortex. RhoGAP15B regulates cluster morphology and is enriched at the leading cell front, where it restrains actomyosin contractility to promote protrusive behavior. Our findings reveal RhoGAP15B as a key spatial RhoA regulator and highlight that patterned RhoGAP and RhoGEF activities are essential for coordinating cortical contraction and protrusion dynamics during collective migration.

## Introduction

Collective cell migration is critical for embryonic development, tissue repair, and supports cancer metastasis (Cheung and Horne-Badovinac, 2025; Pena and Martin, 2024). The directed migration of a multicellular group relies on the collective reorganization of the cytoskeleton in response to external guidance cues. The ability of migrating cells to quickly remodel their cytoskeleton to coordinate protrusive with contractile behaviors is orchestrated by the Rho family GTPases, most notably RhoA, Rac1, and Cdc42, which act as fast molecular switches to regulate actin polymerization, myosin activation, and polarity signaling (Nobes and Hall, 1995; Ridley, 2015; Warner et al., 2019). Thus, understanding how RhoGTPase activities are controlled with spatiotemporal precision is crucial to elucidate the mechanisms that enable efficient collective migration.

Rho GTPases alternate between active (GTP-bound) and inactive (GDP-bound) states through the opposing actions of Rho guanine nucleotide exchange factors (RhoGEFs) and Rho GTPase-activating proteins (RhoGAPs), which respectively promote GTP loading and accelerate GTP hydrolysis. RhoGEF/GAPs are large protein families that comprise 145 members in humans and 46 in Drosophila, and must therefore underlie the dynamic pattern of RhoGTPase activity. RhoGAPs and GEFs have been widely studied during cell division, morphogenesis and tissue repair (di Pietro et al., 2023; Fic et al., 2021; Garcia De Las Bayonas et al., 2019; Jackson et al., 2024; Laurin et al., 2019; Mason et al., 2016; Nakamura et al., 2017; Silver et al., 2019), while a variety of individual RhoGAP/GEFs have also been studied in migrating cells (reviewed in (Lawson and Ridley, 2018)). However, there are no systematic studies addressing how these regulators are deployed during collective cell migration in vivo. These are critical to understand the dynamic patterns of Rho GTPase activity, as they rely on the combinatorial activity of multiple GEFs/GAPs, integrated in feedback loops that are responsive to specific stimulus and self-reinforcing (Bement et al., 2024).

The *Drosophila* border cell cluster provides a powerful system to investigate collective cell migration *in vivo* (Montell et al., 2012). It consists of a pair of non-migratory polar cells surrounded by six to eight migratory BC that delaminate from the anterior follicular epithelium during stage 9 of oogenesis and migrate between nurse cells until reaching the oocyte. The core set of Rho family GTPases controls distinct aspects of BC migration. Rac1 responds to chemokine receptors in leader cells, promoting the formation of actin-rich protrusions that guide cluster movement (Cai et al., 2014; Fernandez-Espartero et al., 2013; Wang et al., 2010), and also promotes follower cell crawling (Campanale et al., 2022). Cdc42 controls the BC cluster apical-basal polarity and cooperates with Rac1 to regulate protrusion actin flows (Wang et al., 2018; Zhou et al., 2022). RhoA and associated myosin activation generate contractile forces at the cortex that sustain the mechanical constraints imposed by nurse cells (Aranjuez et al., 2016), as well as dynamic Myosin flashes that retract ectopic protrusions (Mishra et al., 2019), control the transition between linear to rotational movement (Combedazou et al., 2017) and push the BC nuclei in-between nurse cells (Penfield and Montell, 2023). This knowledge highlights that spatial patterning of Rho GTPase signaling is essential for BC movement, but whereas several GEFs (Vav (Fernandez-Espartero et al., 2013; Poukkula et al., 2011), Mbc (Bianco et al., 2007), Cdep (Campanale et al., 2022), sif (Wang et al., 2018)) and a GAP (RhoGAP18B, (Lei et al., 2023)) have been implicated in Rac1 regulation, it remain unknown how RhoA and Cdc42 activities are regulated during BC migration.

Here, we performed a family-wide localization-function screen to assess the role of RhoGAPs and RhoGEFs during Drosophila BC migration. This approach identified several previously uncharacterized RhoGAP/GEFs as putative cytoskeleton regulators, including the conserved RhoA inhibitor RhoGAP15B. RhoGAP15B plays a localized role at the cortex of the leading cell, where it restrains RhoA-dependent contractility to stabilize front-directed protrusions. Importantly, precisely tuned RhoA inhibition is necessary to coordinate protrusive activity with actomyosin contraction as both loss and overexpression (OE) of RhoGAP15B affect protrusion dynamics and impair movement. Our findings uncover a new layer of spatiotemporal control in collective cell migration, suggesting that localized relaxation at the leading front, coordinated by RhoGAP15B, organizes the balance between contractile and protrusive behaviors.

## Results and Discussion

### Family-level analysis of GEF/GAP localization and function in BC migration

To investigate the regulatory network that orchestrates the spatiotemporal of RhoGTPases during BC migration we analyzed the subcellular localization of *Drosophila* RhoGEFs and RhoGAPs. We used a library comprising all 26 *Drosophila* RhoGEFs, and 22 RhoGAPs, each endogenously tagged with GFP using CRISPR/Cas9 (di Pietro et al., 2023), and co-stained egg chambers for F-actin to visualize BC cluster morphology. From this analysis, we identified 8 RhoGAPs and 9 RhoGEFs that are strongly expressed in BCs (overview in Table S1, Fig. 1A). This set included Cdep and Vav, which have been previously implicated in BC migration ((Campanale et al., 2022; Fernandez-Espartero et al., 2013; Poukkula et al., 2011)), but most identified RhoGAPs and RhoGEFs play yet uncharacterized roles in BC migration. Furthermore, this analysis revealed a complex localization pattern, including polar cell-specific or BC nuclear accumulation (Fig. 1B) and broader cytoplasmic distribution (Fig. 1E). Interestingly, many RhoGEFs and RhoGAPs displayed distinctive cortical enrichment patterns that could be associated with cytoskeleton regulation. Their localization suggested potential roles within leading cell protrusions (Sos; RhoGAP15B; Pbl - Fig. 1B, 1C); in the regulation of actomyosin flashes in the cortex of follower cells (cdGAPr, CG43658; Conu; RtGEF - Fig. 1E), and in intercellular coupling within the cluster, where several proteins accumulated at contacts between polar and BCs (e.g. Zir, Trio, Graf, RhoGAP19D, arrows in Fig.1D). In addition, several RhoGEFs (RtGEF, Cysts, Pbl) and RhoGAPs (RhoGAP15B, RhoGAP19D, Graf, Conu) displayed local enrichment at the BC cortex in the detachment of BCs from the epithelial monolayer, and during the neolamination step, when clusters reconnect with the epithelium (Fig. S1). Together, this comprehensive analysis unveils a set of RhoGEF/GAPs localization patterns that may contribute to distinct regulatory steps of BC migration.

**Figure 1).**
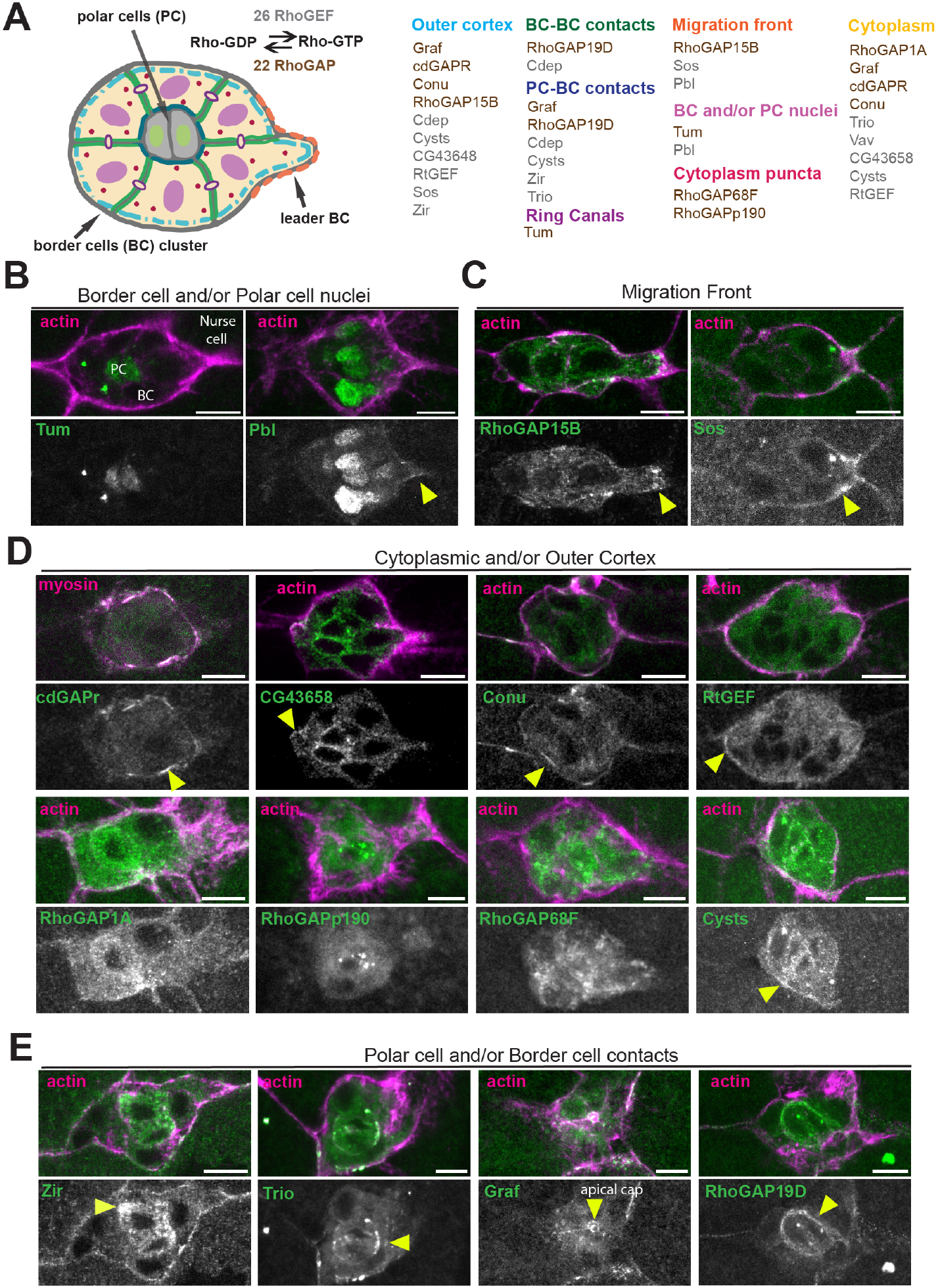
Localization of RhoGEF and RhoGAPs during BC migration. **(A)** Schematic representation of the subcellular localization of endogenously GFP-tagged RhoGAPs and RhoGEFs detected during border cell migration. **(B-E)** Confocal images of RhoGEF/GAPs with distinct localization patterns: nuclear enrichment in border cells and/or polar cells (B); accumulation at the leading cell front (C, yellow arrowhead in B, C); with cytoplasmic distribution with local enrichment at the outer cortex (D, arrowheads) and enrichment at the polar cell – border cell contacts (E, arrowheads). Egg chambers were stained for actin or co-expressed Sqh:3xmKate2 (magenta) to mark the border cell cortex. GFP signals are also shown separately (bottom panels). Scale bars, 10 μm.

Having identified 15 GAP/GEFs that remain uncharacterized in BC migration, we sought to investigate their functional significance by a targeted RNAi-based screen. To quantify migration defects, we measured the relative position of the BC cluster relative to the oocyte position at stage 10 of oogenesis. The mild protein depletions often achieved by RNAi-mediated knockdown, complicates the systematic evaluation of the functional importance of individual RhoGTPase regulators. We therefore implemented a genetic strategy outlined below to assess BC migration defects while ensuring robust validation of protein depletion (Fig. S2A). First, we used drivers that initiate Gal4 expression in the follicular epithelium at early stages of oogenesis, namely *GR1-GAL4* or alternatively *tj-GAL4* (Wittes and Schupbach, 2019) to express UAS-RNAi lines. By extending the RNAi expression period (~ 36 h of RNAi expression from stage 4 to stage 10 of oogenesis in the case of GR1-GAL4 and an even longer period for *tj-GAL4*), it promoted the efficient knockdown of highly stable proteins. Second, we combined RNAi-expression in the context of the corresponding GFP-tagged RhoGAPs/RhoGEFs to track *in situ* their depletion. To further overcome the challenge of detecting weak fluorescence signals, we further included a GFP-clustering module based on light-induced CRY2/CIBN multimerization (Qin et al., 2017)). This approach amplifies GFP signals, enabling reliable protein detection even under low expression conditions. Using this system, we compared the migration rate of each RNAi condition to its respective protein clustering control.

RNAi-mediated depletion of 6 RhoGEFs (Cysts, CG43658, Zir, Sos, Pbl and Vav (positive control)) and 5 RhoGAPs (Conu, Graf, RhoGAP15B, Tum and cdGAPr) resulted in significant BC migration defects (Fig. 2A-2C, Fig. S2B-S2J and Table S1). However, despite the strong protein depletion achieved for most RhoGAPs/RhoGEFs analysed (Table S1 and Fig. S2), none fully disrupted BC movement. These findings suggest that many RhoGAPs and RhoGEFs contribute to BC migration, but their roles in regulated detachment and movement are at least partly redundant. Compensatory mechanisms may also ensure successful migration of the BC cluster, highlighting the robustness of collective cell behaviors. Further experiments with multiple genetic perturbations are required to confirm the contribution of each RhoGAP/GEF. Here, we focused on RhoGAP15B because depletion resulted in one of the most penetrant defects (Fig. 2B and Fig. 2C). In addition, RhoGAP15B and its human orthologues (ARAP1/3) interact with the RhoA type of GTPases to control Rho-GTP hydrolysis (Bao et al., 2016; Imran Alsous et al., 2021), and unlike Rac1 regulation, the spatial regulation of RhoA during BC migration remains unexplored.

**Figure 2).**
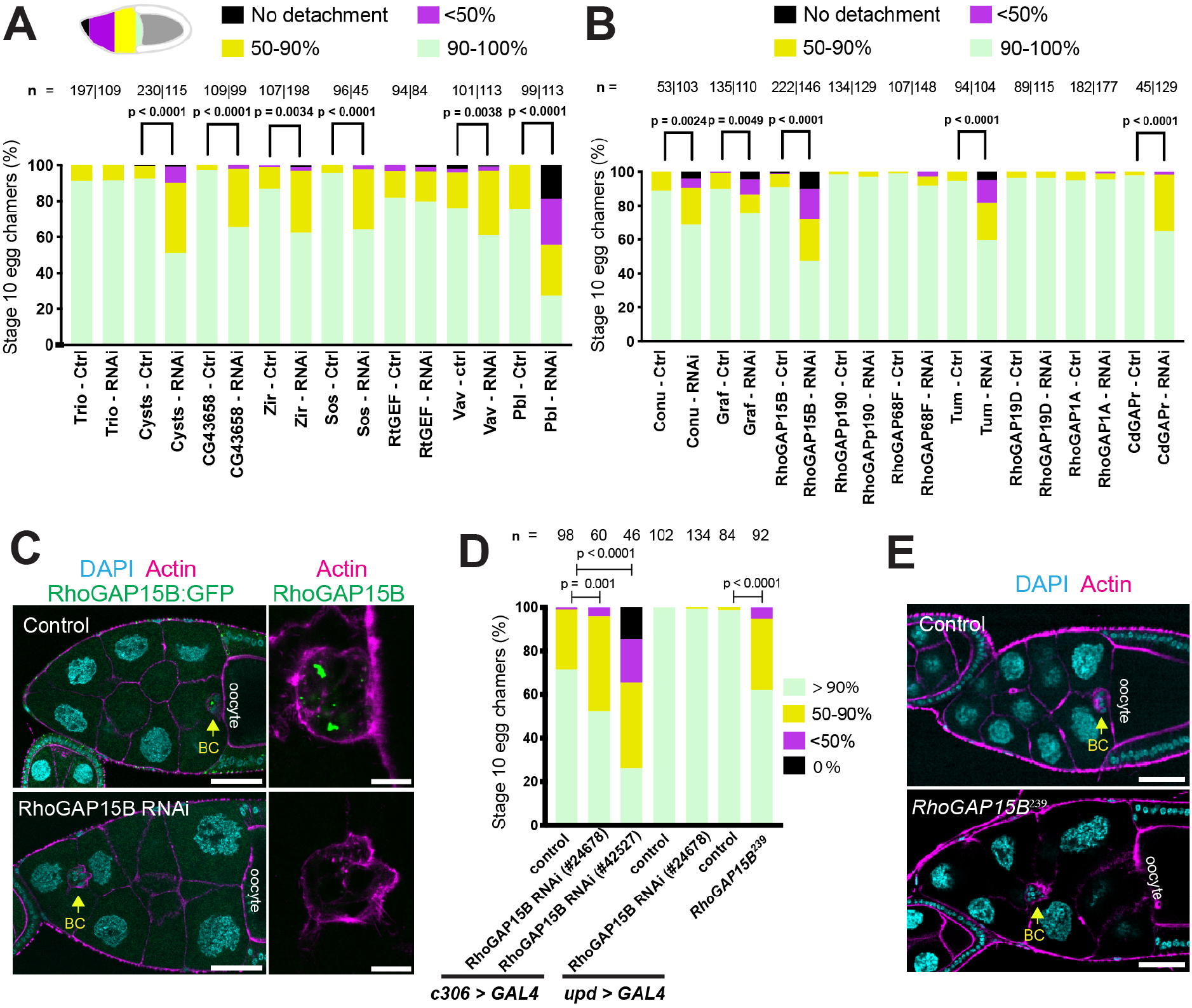
Identification of RhoGAP15B in a screen for border cell migration defects. (**A, B**) Frequency distribution of BC migration delay in RNAi-mediated depletion of RhoGEFs (A) or RhoGAPs (B) and the respective controls. Detailed screen genotypes are provided in Table S4. Mann-Whitney tests for statistical significance were performed on raw migration ratio data. n indicates the number of stage 10 egg chambers analysed. See also Figure S2. (**C**) Representative confocal image showing delayed BC migration upon RhoGAP15B depletion in the RNAi screen conditions. Egg chambers expressing RhoGAP15B:GFP (green) were stained for DAPI (cyan) and F-actin (magenta). Enlarged views of the BC cluster confirm RhoGAP15B depletion. Arrows indicate BC position. Scale bars: wide view - 50 μm; close up - 10 μm. (**D**) Quantification of BC migration delay after depletion of RhoGAP15B using distinct RNAi lines driven by a BC-specific driver (c306-Gal4) or polar cell-specific driver of expression (upd-Gal4), and in RhoGAP15B239 mutants. Mann-Whitney tests for statistical significance were performed on raw migration ratio data. n indicates the number of stage 10 egg chambers analysed. (**E**) Confocal images showing delayed BC migration in RhoGAP15B239 mutant egg chambers. Egg chambers were stained for DAPI (cyan) and F-actin (magenta). Scale bars: 50 μm.

### Balanced RhoGAP15B activity is required for efficient BC migration

Recent work has shown that RhoGAP15B regulates contractile actomyosin waves in the germline during nurse cell dumping in late Drosophila oogenesis (Imran Alsous et al., 2021; Jackson et al., 2024), but its functions in migrating cells have not been described. To corroborate the BC function of RhoGAP15B, we used two other UAS-driven RNAi lines (VDRC #24678 and BDSC #42527 (TRiP.HMJ02093)), and further restricted RNAi expression to the BC cluster (c306-GAL4) or specifically to polar cells (upd-GAL4). In addition, we analyzed the BC migration index in null RhoGAP15B239 mutant lines generated by deleting most of the coding region of all RhoGAP15B isoforms (N239 to K1552 of isoform RC). All genetic manipulations that perturbed RhoGAP15B BC function disrupted the migration of the cluster, whereas depletion of RhoGAP15B only in polar cells did not cause migration defects (Fig. 2C-E) indicating that RhoGAP15B is specifically necessary in BCs. To further investigate which migratory features were impaired, we tracked BC migration by time-lapse imaging of BCs expressing an actin cytoskeleton marker LifeAct:GFP (Fig. 3A-C). Both mutant egg chambers and BC clusters expressing RhoGAP15B RNAi under the control of c306-GAL4 fully delaminate from the follicular epithelium, with only a few cases of delayed delamination (Fig. 3A; wild-type delamination occurs before oocyte length reaches 70 μm (Inaki et al., 2022)). Quantification of BC migration speed revealed that both mutant and RNAi-depleted clusters are consistently slower than controls (Fig. 3B and 3C, Movie S1), suggesting that RhoGAP15B is primarily required for efficient cluster migration.

**Figure 3).**
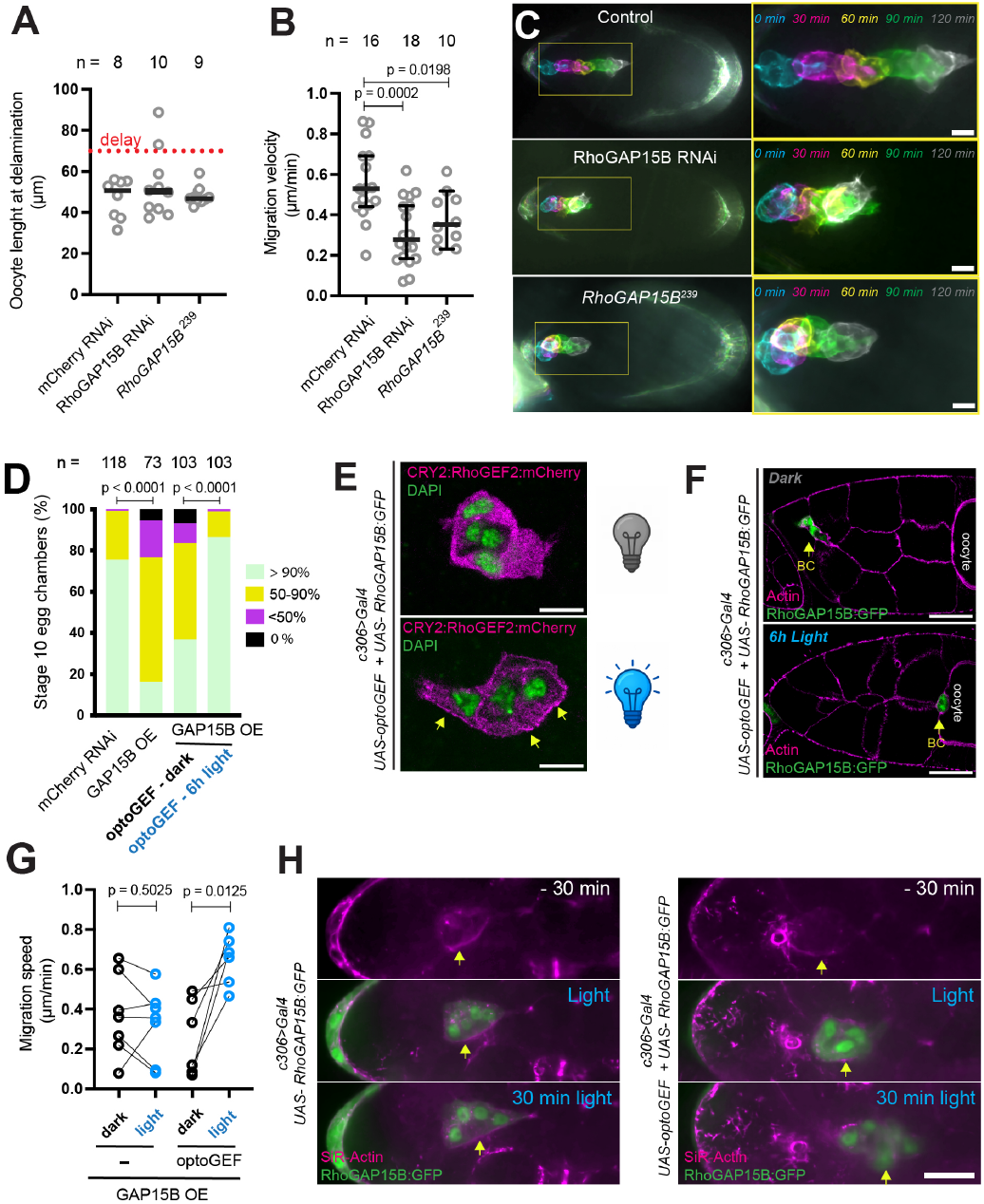
Balanced RhoGAP15B activity is required for efficient BC migration. **(A)** Oocyte length at delamination, used as a proxy of delay in delamination. A threshold of 70 μm corresponds to the maximum wild-type length at delamination as defined previously ((Inaki et al., 2022). **(B)** Quantification of BC cluster migration velocity in egg chambers cultured ex-vivo. Each dot represents one cluster. Error bars show the 95% confidence interval. Statistical significance was assessed using one-way ANOVA. **(C)** Time color-coded images from 2-hour timelapse series of stage 9 egg chambers imaged since delamination. *slbo-LifeAct:GFP* marks BCs in control, RhoGAP15B RNAi (VDRC #24678) and *RhoGAP15B*^*239*^ mutant backgrounds. Close ups of the migrating cluster are shown on the right. Scale bars: 10 μm. (**D)** Quantification of BC migration delay upon OE of UAS-RhoGAP15B:GFP driven by *c306-Gal4*, and after co-expressing the optoGEF optogenetic module (UAS-pmCIBN and UAS-RhoGEF2:CRY2:mCherry) under dark (cytosolic RhoGEF2) or after 6 hour of blue light exposure (membrane-recruited RhoGEF2). Statistical significance was assessed using Mann– Whitney tests on raw migration ratio data. *n* indicates the number of stage 10 egg chambers. **(E and F)** Confocal images of BC co-expressing RhoGAP15:GFP and the optoGEF module in dark (top) and blue light (botton) conditions. Light exposure recruits RhoGEF2:CRY2:mCherry (magenta, arrows) to the membrane (E) and rescues migration defects caused by RhoGAP15B:GFP OE (F, arrows). Egg chambers were stained for nuclei (DAPI, green in E) and actin (magenta in F). Scale bars: wide view, 50 μm; close-up, 10 μm. **(G)** Quantification of migration speed in RhoGAP15B:GFP-overexpressing clusters in the absence or presence of optoGEF imaged during consecutive 30 min periods under dark and light conditions. Each dot represents one cluster; statistical significance was determined using parametric paired *t*-tests. **(H)** Representative time-lapse images showing BC migration prior to blue-light exposure, at the onset of light activation and 30 min after. The cell cortex was visualized using SiR-Actin (magenta). Scale bar, 20 μm.

RhoGAP15B OE also caused BC migration defects (Fig. 3D), suggesting that proper BC migration requires balanced GAP and GEF activities. To test this hypothesis, we asked if restoring the increasing cortical GEF availability could restore the migration of RhoGAP15B overexpressing cells. We used an optogenetic system to trigger membrane recruitment of RhoGEF2 (RhoGEF2:CRY2:mCherry) via the blue-light dependent interaction with CIBN:CAAX (optoGEF (Izquierdo et al., 2018)). Living flies co-expressing UAS-RhoGAP15B:GFP and the UAS-driven optoGEF in BC clusters were exposed to blue light for six hours prior to fixation, allowing RhoGEF2 cortical recruitment during the migration period (Fig. 3E). OptoRhoGEF activation significantly rescued migration compared to UAS-RhoGAP15B:GFP overexpressing clusters kept in the dark (Fig. 3D, 3F). To directly test the effect of acute RhoGEF2 activation on the migratory ability of RhoGAP15B-overexpressing clusters, we compared the migration speed of delaminated BC clusters in the 30 minutes before and after light activation. While clusters overexpressing only UAS-RhoGAP15B:GFP maintain slow migration speed, clusters co-expressing the optoGEF exhibited increased motility upon blue-light exposure (Fig. 3G, 3H and Movie S2). Together, these findings indicate that efficient BC migration is governed by the balanced activity of RhoGAP15B and counteracting RhoGEFs, ensuring dynamic cycles of RhoA GTPase activity.

### RhoGAP15B promotes the protrusive activity of leading cells

RhoGAP15B shows local subcellular enrichment associated with the distinct axes of BC polarity (apical-basal, outside-inside, front-rear) (Wang et al., 2018). Similarly to its accumulation in the apical side of follicle cells, RhoGAP15B is also enriched in the apical cap of polar cells in migrating clusters (Fig. S3A). However, *RhoGAP15B* BC mutant clones do not show defects in adherens junction distribution or apical-basal polarity (Fig. S3B-S3E). RhoGAP15B:GFP also displays local enrichment in patches at the outer cortex of the BC cluster (Fig. 4A and S1A) and accumulates at the protrusive leading cell, adjacent to protrusive filamentous actin (Fig. 1C; 4B, S1A). As a first approach to investigate the impact of RhoGAP15B on the BC cytoskeleton, we tested if RhoGAP15B loss of function reproduced cell shape changes associated with Rho activation. Overexpression of a constitutively active version of RhoA (RhoA-CA (RhoA^V14^)) induces BC rounding (Fig. S4A), a hallmark of cortical hypercontractility, as previously described (Aranjuez et al., 2016; Combedazou et al., 2017). Similarly, BC clusters from RhoGAP15B mutant egg chambers show an increased number of rounded cells (Fig. S4A and Movie S3), indicating that RhoGAP15B restrains actomyosin contractility.

**Figure 4.**
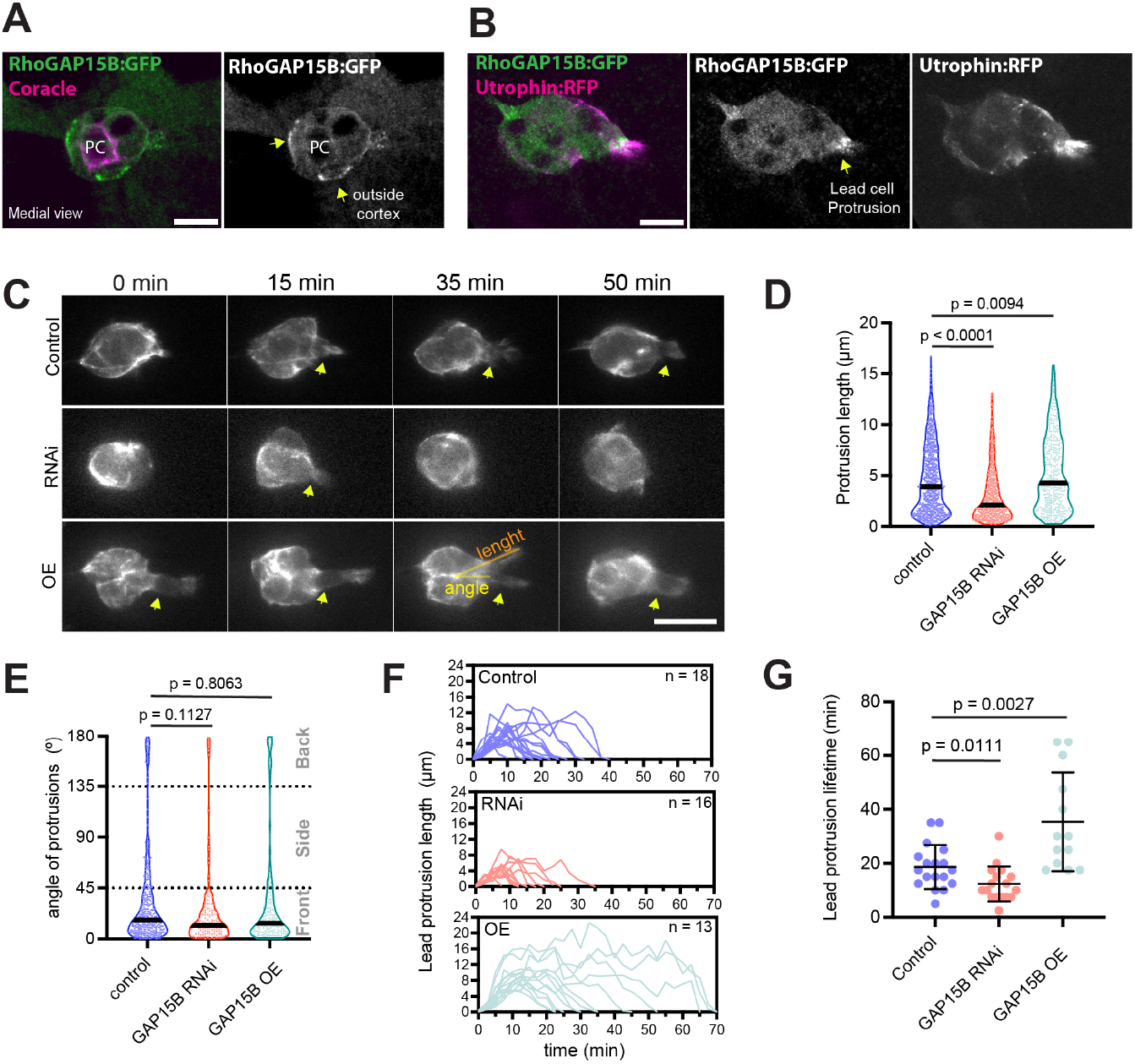
RhoGAP15B promotes protrusive activity in leading BCs. **(A and B)** Confocal images showing the localized accumulation of RhoGAP15B:GFP at the outer BC cortex (A, arrows) and adjacent to actin filaments at the leading protrusion (B). Coracle labels polar cell (PC) septate junctions in A, and the F-actin reporter Utrophin:RFP marks the leading protrusion in B. **(C)** Confocal timelapse images of migrating BCs in stage 9 egg chambers expressing slbo-LifeAct:GFP in control, RhoGAP15B RNAi and RhoGAP15B OE. Orange line depicts the ROI to measure protrusion length and the yellow lines depict how the protrusion angle relative to the anterior-posterior axis was accessed. Arrows mark lead protrusions. Scale bars: 20 μm. **(D and E)** Quantification of protrusion length (D) and main protrusion angle (> 4 μm) (E) during BC migration in control (n = 16), RhoGAP15B-RNAi (n = 15), and RhoGAP15B-OE (n = 8) clusters. Violin plot show individual values measured from 12 frames of 1h movies. Mann–Whitney tests were used to determine statistical significance.**(F and G)** Lead protrusion dynamics (F) and lifetime (G) from the initial extension to full retraction in control (n=18), RhoGAP15B RNAi (n=16), and OE of RhoGAP15B (n=13). Statistical significance was assessed using Mann-Whitney tests.

Intriguingly, both RhoA-CA expression and RhoGAP15B mutation resulted in a marked reduction of front-directed protrusions detected in fixed BC clusters (Fig. S4B). To further elucidate the role of RhoGAP15B in regulating protrusion dynamics, we performed live imaging using the actin reporter LifeAct:GFP in BCs perturbed by either RhoGAP15B RNAi or RhoGAP15B OE (Fig. 4C). In timelapse movies, RhoGAP15B RNAi reduced the length of protrusions, whereas RhoGAP15B OE had the opposite effect (Fig. 4C-4D, Movie S4). Notably, main protrusions (> 4 μm) remained largely confined to the leading cells, as few protrusions were detected at the side or the back in RhoGAP15B perturbations (Fig. 4E). To quantify protrusion dynamics, we tracked the length of lead protrusions over time. RhoGAP15B RNAi reduced, whereas RhoGAP15B OE enhanced protrusion persistence (Fig 4F and 4G). Overall, this result reinforces the significance of RhoGAP15B in BC migration, controlling protrusive behaviour to define the length and temporal stability of front-directed protrusions.

### Localized Rho inhibition restricts actomyosin contraction to stabilize lead protrusions

Regulation of actomyosin contractility modulates the protrusive behavior of BCs (Lamb et al., 2021; Mishra et al., 2019; Plutoni et al., 2019). We therefore hypothesized that RhoGAP15B could regulate protrusion dynamics by regulating RhoA-mediated actomyosin contractility. We first asked whether the local front enrichment of RhoGAP15B could be associated with an asymmetry of Rho activity in the BC cluster. Consistent with this, live imaging Förster Resonance Energy Transfer (FRET) analysis using a Rho activity reporter shows that Rho activity is lower in the lead protrusion (Fig. 5A and 5B). In contrast, BC clusters depleted of RhoGAP15B do not show asymmetric Rho activity and exhibit increased in Rho activity (Fig 5A and 5C), supporting RhoGAP15B role as a RhoA inhibitor during BC migration. To determine the effect of RhoGAP15B RNAi on myosin II dynamics, we co-imaged LifeAct together with fluorescently tagged myosin light chain (Sqh:mScarlet). RhoGAP15B RNAi did not significantly affect the frequency of cortical myosin flashes or the overall cortical myosin levels (Fig. 5D-5F). In control movies, myosin flashes are frequently observed behind the leading cell cortex and in the cortex of follower cells (Fig. 5G and 5H). By contrast, RhoGAP15B depletion shifted myosin towards the leading front, where myosin pulses are associated with the fast retraction of protrusions (Fig. 5G and 5H and Movie S5). Quantification of myosin distribution during protrusive periods confirmed front myosin enrichment along the front-rear axis in RhoGAP15B RNAi clusters (Fig. 5H). Thus, our results suggest that RhoGAP15B suppresses actomyosin contractility in the leading cell front to sustain protrusion extension.

**Figure 5.**
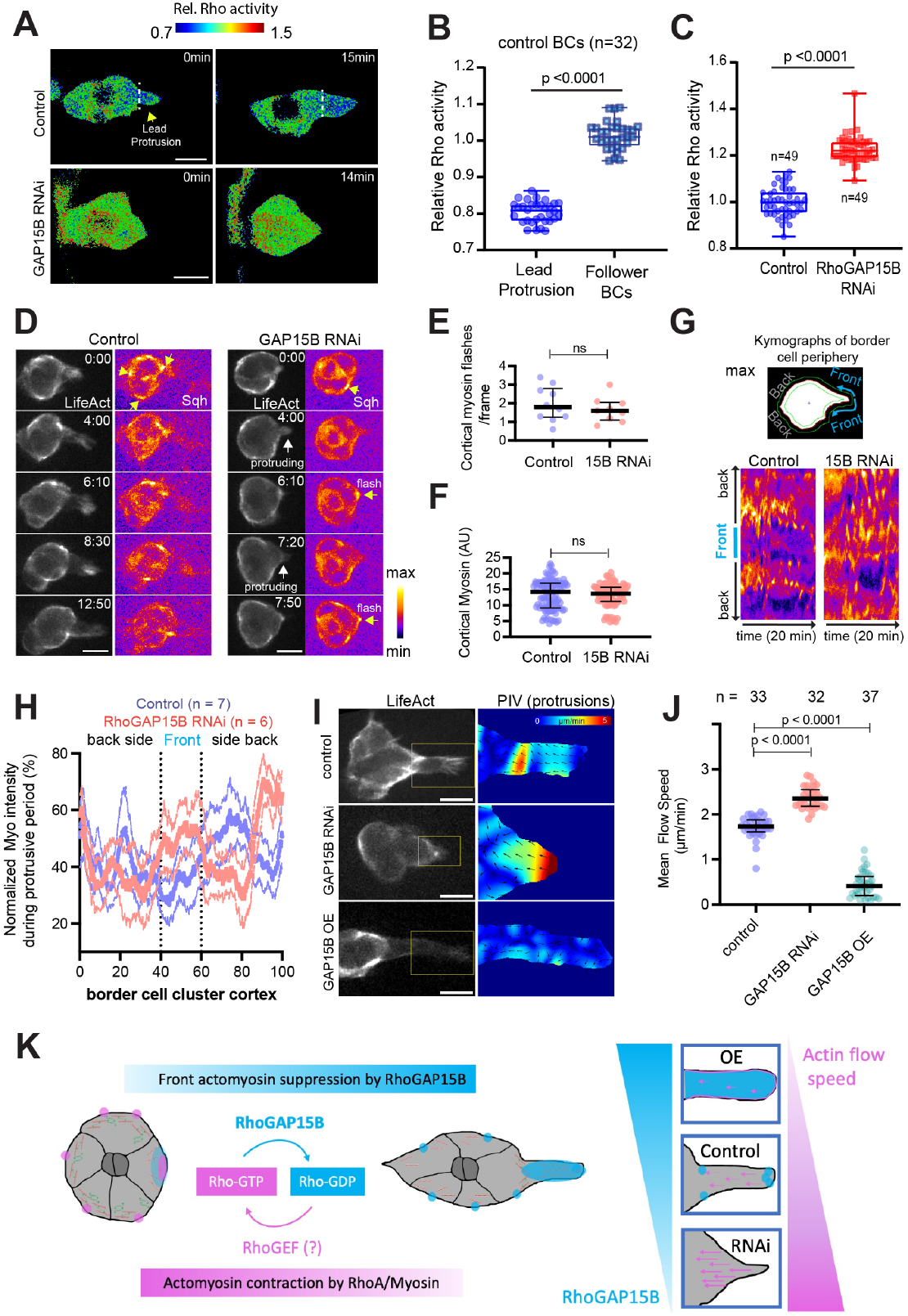
Localized Rho inhibition stabilizes protrusions by limiting actomyosin contractility. **(A)** Representative images of Rho activity (RhoFRET) in control (co-expressing UAS-RhoFRET and UAS-mCherry RNAi driven by c306-Gal4) and in RhoGAP15B RNAi (co-expressing UAS-RhoFRET and UAS-RhoGAP15B RNAi driven by c306-Gal4). **(B)** Quantification of Rho activity in leader cell protrusions versus follower BCs in control egg chambers. Each dot represents one BC cluster. Statistical significance was assessed using Student’s t-test. Boxplots indicate medians, 25th and 75th percentiles (boxes), minimum and maximum values (whiskers), and all data points (dots). **(C)** Quantification of Rho activity in control and *RhoGAP15B* RNAi egg chambers. Each dot represents one BC cluster. Statistical significance was assessed using Student’s *t*-test. Boxplots indicate medians, 25th and 75th percentiles (boxes), minimum and maximum values (whiskers), and all data points (dots). **(D)** Time-lapse images comparing local myosin enrichment and protrusive behavior in control or RhoGAP15B RNAi. *slbo-LifeAct:GFP* marks actin (gray) and *sqh-mScarlet* marks non-muscle myosin II (fire LUT). Arrows mark cortical myosin flashes. Note front myosin flashes retracting protrusions in RNAi condition. Scale bar, 10 μm. **(E, F)** Quantification of cortical myosin flashes (E) and mean cortical myosin intensity (F) in control (*n* = 10) and RhoGAP15B-RNAi (*n* = 9) clusters. Scatter plots show mean number of cortical myosin flashes per movie in (E) and individual values in (F). Statistical significance was assessed using Mann–Whitney tests. (**G**) Kymographs of myosin intensity along the BC cortex (obtained from the time-lapse movie presented in H) in control and RhoGAP15B RNAi depleted clusters. (**H**) Normalized plot profiles of myosin intensity along the BC cortex during protrusive phases BCs in migrating control (n = 7) and RhoGAP15B RNAi (n=6) BC clusters. **(I, J)** PIV analysis (I) and quantification of mean flow speed (J) of F-actin flows in the leading protrusion of control, RhoGAP15B RNAi, and RhoGAP15B OE BC clusters expressing LifeAct-GFP for F-actin signal (I, left panel). PIV analysis (I, right panel) performed on the LifeAct-GFP signals to highlight the direction and magnitude of actin flows. Yellow boxes mark the regions of interest used for PIV. Each dot in (J) represents one leading protrusion. Lines indicate median and interquartile range. *P* values were determined by the Kruskal–Wallis test followed by Dunn’s multiple comparisons test. Scale bar (J), 5 μm. **(K)** Model illustrating the role of RhoGAP15B to promote protrusive activity by locally restricting RhoA-mediated actomyosin contraction at the cluster front. RhoGAP15B-mediated inhibition of Rho reduces actin retrograde flows, maintaining actin available for polymerization at the protrusion tip. This spatial regulation ensures the persistence of front-directed protrusions while preserving overall cluster shape.

Recent work identified retrograde actin flows initiating near the leading protrusion tip, which coordinate protrusive with contractile properties to ensure BC migration efficiency (Zhou et al., 2022). Given the role of RhoGAP15B in regulating protrusive behavior, we applied Particle Image Velocimetry (PIV) to LifeAct:GFP fluorescence to assess if RhoGAP15B perturbations alter actin flows in the protrusion region. RhoGAP15B depletion significantly increased, while OE reduced, the mean speed of actin flows (Fig. 5I and 5J). These findings are consistent with the fact that elevated myosin contraction accelerates retrograde actin flows, whereas myosin suppression reduces them (Cai et al., 2006; Qian et al., 2024; Yang et al., 2012; Yao et al., 2023; Yolland et al., 2019). Altogether, our results suggest that RhoGAP15B-mediated RhoA inhibition is critical to confine contractility and balance retrograde flow, preventing an excessive flow towards the protrusion neck that can destabilize the lead protrusions (Fig. 5K).

## Conclusion

Collective cell migration arises from the precise spatial and temporal coordination of Rho GTPase signaling. Our analysis of RhoGEF and RhoGAP distributions in *Drosophila* BCs reveals a complex network of Rho GTPase regulators acting in distinct cortical domains. While previous studies have characterized Rac1 activators within the BC cluster (Bianco et al., 2007; Campanale et al., 2022; Fernandez-Espartero et al., 2013; Poukkula et al., 2011), our work extends the mechanistic understanding of RhoGTPase regulation by identifying RhoGAP15B as a critical RhoA inhibitor required to sustain protrusion formation and stabilization. In addition, the localization of other hits from the RNAi screen, including the RhoGEFs Sos, Cysts and Pebble, and the RhoGAPs Conu and Graf, pinpoint new putative regulators the dynamic RhoGTPase-dependent cytoskeleton rearrangements that drive BC migration.

While RhoGAP15B contributes to the overall morphology of the BC cluster, its primary function is to locally restrain actomyosin contractility to ensure protrusion persistence in the leading cell. This is consistent with previous findings that transient Myosin II accumulation promotes protrusion retraction during BC migration (Mishra et al., 2019). RhoA activation has similarly been associated with protrusion retraction in diverse cellular contexts (Azoitei et al., 2019; Hu et al., 2022; Martin et al., 2016), underscoring the importance of identifying localized RhoA inhibitors regulating protrusive behaviour. RhoFRET analysis revealed a spatial compartmentalization of RhoA activity within BC clusters, with lower activity at the leader cell protrusion. This is complementary to front-localized Rac1 activity (Wang et al., 2010), and consistent with a functional antagonism between these Rho GTPases. Our findings indicate that RhoGAP15B suppresses retrograde actin flows that would otherwise deplete the actin pool available for polymerization at the protrusion tip, suggesting that RhoGAP15B-mediated RhoA inactivation could also promote Rac1-driven branched actin polymerization. Nevertheless, while RhoGAP15B inactivates RhoA in leading cells, active RhoA must cooperate with a distinct Rac1 pool to sustain supracellular actin cables that regulate morphology of the peripheral cluster cortex and mediate intercellular communication (Wang et al., 2020; Zhou et al., 2022). These cortical actin networks also act in concert with the Mishapen-Moesin pathway to regulate contractility and restrain ectopic protrusion formation (Plutoni et al., 2019). Thus, RhoGAP15B must function as a spatial regulator that balances the protrusive activity of leader cells while maintaining the collective contractility of the BC cluster.

OE of RhoGAP15B also perturbs BC migration, an effect rescued by increasing RhoGEF cortical availability. This finding illustrates the importance of integrating RhoGEF and RhoGAP-mediated feedbacks to maintain dynamic RhoA activation-inhibition, thereby coordinating cycles of protrusion extension and retraction, as reported in the zebrafish posterior lateral line primordium (Qian et al., 2024). An important future challenge will be resolving whether similar pulsatile RhoA dynamics occur during BC migration, while identifying the antagonistic RhoGEF that promotes protrusion retraction. Notably, RhoGAP15B was shown to act in concert with the RhoGEF Pbl to generate actomyosin waves during nurse cell dumping (Jackson et al., 2024), suggesting that this GAP-GEF pair could also underlie contraction-relaxation cycles in migrating contexts. Interestingly, the mammalian RhoGAP15B orthologue, ARAP3, suppresses RhoA activity through localized cortical accumulation to regulate lamellipodia formation in endothelial cells (Krugmann et al., 2006). Hence, spatial restriction of Rho actomyosin contractility by RhoGAP15B/ARAP3 may represent a conserved mechanism regulating protrusion dynamics across diverse cellular contexts.

## Materials and Methods

The list of reagents used in this study is found in table S2.

### Drosophila maintenance and genetics

Fly stocks and genetic crosses were raised in standard fly media (cornmeal/agar/molasses/yeast) at 18°C or 25°C, with 60% humidity and 12h/12h dark light cycle, unless stated otherwise. The fly lines used throughout this study are listed in Table S3. A detailed list of the fly genotypes for each experiment can be found in Table S4. We used tissue specific Gal4 lines to drive the expression of UAS constructs in the Drosophila BCs (slbo-Gal4 or c306-Gal4), polar cells (upd-Gal4), follicular epithelium (tj-GAL4 or GR1-Gal4).

### In vivo genetic screen with depletion validation system

The RhoGAP/GEF RNAi lines were obtained from Drosophila Bloomington Stock Center (DBSC) or Vienna Drosophila Resource Center (VDRC), and all genotypes are listed in Table S4. In general, flies containing: 1) a follicular epithelium specific driver (tj-Gal4 or GR1-Gal4); 2) UAS-GAP or UAS-GEF RNAi; 3) endogenously expressed GFP-tagged GAP or GEF (di Pietro et al., 2023); 4) UAS-driven LARIAT to activate clustering (Qin et al., 2017). All crosses were performed at 18°C and 0 to 4 days after hatching, adult offspring was transferred to 29°C for 3 days prior to dissection to boost the efficiency of RNAi depletion. The period of incubation at high temperature was increased if depletion was inefficient. At least 2 fully independent experiments were performed for each RNAi line used and, for each of these independent experiments, a minimum of 25 stage 10 egg chambers (randomly selected from a pool of 8-10 dissected animals) were analyzed.

### Generation of UAS-RhoGAP15B:GFP and UAS-RhoGAP15B:HA transgenic lines

RhoGAP15B coding sequence was amplified by PCR from rhoGAP15B cDNA (clone SD08167 (Drosophila Genomics Resource Center (DGRC) Stock 5443;) using the primers: 5’-GCTGCTCCATTTACAATGGATCTGGACAGGCGGCA-3’ / 5’-AGCGCGTCCACCTTTCTTGATGATGATCTCCGCCG-3’. PCR amplifications were performed with Phusion Polymerase (Thermo Fisher). These primers created overlapping ends to the pENTR entry vector so that pENTR-RhoGAP15B could be generated using the FastCloning (Li et al., 2011) strategy. The Gateway cloning system (Thermo fisher scientific) was used to generate UAS-RhoGAP15B:GFP and UAS-RhoGAP15b:HA plasmids. LR Clonase II (Thermo Fisher) reactions recombined the RhoGAP15B sequence from pENTR-RhoGAP15B into pUASt-attB-WG (Saverio Brogna, University of Birmingham, UK) or into the pUASg-HA-attB (Bischof et al., 2013) destination vectors. UAS-RhoGAP15B:GFP and UAS-RhoGAP15b:HA transgenes were then inserted into the M{3xP3-RFP.attP)ZH-86Fb landing site on chromosome III (BL stock: 24749) via PhiC1 site-specific transgenesis (CONGENTO, Consortium for Genetically tractable organisms, Portugal)

### Generation of RhoGAP15B239 mutant

The RhoGAP15239 mutant was generated as described in (di Pietro et al., 2023), by deleting a 5712 bp genomic region within the RhoGAP15B locus (ChrX: 16,950,512 – 16,956,223). This deletion removes amino acids N239– K1552 of the RhoGAP15B-RC isoform (1552 aa) and thus disrupts all reported RhoGAP15B isoforms. The strategy consists of a double Homology-Directed Repair (HDR) where homology regions were assembled into pCRISPR-del to generate the donor constructs for HDR. The two homology regions (HR1 and HR2) were designed for RhoGAP15B locus were: HR1 consisted of a left homology arm amplified using: Forward primer: 5′-CCGGGCTAATTATGGGGTGTCGCCCTTCGCAACAGTCCTCCGGCAGCCAAG-3′ (homology to 16,949,541– 16,949,561 bp) and reverse primer: 5′-ACTCAAAGGTTACCCCAGTTGGGGCACTACCGACCGCACAAAACTCAGATTTCG-3′ (homology to 16,950,489–16,950,512 bp). HR2 consisted of a right homology arm amplified using: Forward primer: 5′-ACTCAAAGGTTACCCCAGTTGGGGCACTACCGCAAGCACACTTGAATATCACTAAAATG-3′ (homology to 16,956,223–16,956,251 bp) and reverse primer: 5′ GCCCTTGAACTCGATTGACGCTCTTCTGTACCCCAAGCGGCAACTGTAACC-3′ (homology to 16,957,224– 16,957,244 bp). Corresponding guide RNA sequences were cloned into pCFD3-dU6:3gRNA (Port et al., 2014): The Cas9 target sites flanking HR1 were: N-terminal sgRNA: forward 5′-GTCGTCGGTAGGGAATTGTGTAT-3′ (targeting 16,950,533–16,950,547 bp) and reverse 5′-GTCGTGCTTGCGTTTTTATTGGG-3′ (targeting 16,950,537–16,950,551 bp) and C-terminal sgRNA: forward 5′-GTCGTCGGTAGGGAATTGTGTAT-3′, and reverse 5′ AAACCCCAATAAAAACGCAAGCA-3′ (targeting 16,956,212–16,956,230 bp), while the Cas9 target sites flanking HR2 were: N-terminal sgRNA: forward 5′ GTCGTGTATAGGAAAGGTATTGTC-3′, and reverse 5′-AAACGACAATACCTTTCCTATACA-3′ (targeting 16,950,547–16,950,566 bp) and C-terminal sgRNA: forward 5′-GTCGCGAAAAATTGGGTCAGATAA-3′, and reverse 5′-AAACTTATCTGACCCAATTTTTCG-3′ (targeting 16,956,095–16,956,114 bp). Plasmids were sent and injected by BestGene Inc. into w Drosophila melanogaster embryos expressing germline Cas9 (nos-Cas9). Candidate transformants were identified by mini-white marker expression in the eyes. Putative RhoGAP15B mutant lines were validated by PCR using primers flanking the deleted region.

### Fixation and staining of egg chambers

Drosophila ovaries were dissected in PBS with 0.05% Tween 20 (Sigma-Aldrich) and fixed using a 4% paraformaldehyde solution (prepared in PBS with 0.2% Tween 20 (Sigma-Aldrich)) for 30 minutes in an orbital shaker at room temperature. After washing three times for 10 minutes with PBT (PBS with 0.05% Tween 20), samples were mounted with Vectashield Mounting Medium with DAPI (Vector Laboratories). For antibody staining, after the post-fixation washes, egg chambers were washed twice for 10 min with “NP-40 Block” as described previously. “NP-40 Block” is an optimized buffer solution containing 50 mM Tris-HCl, pH 7.4, 15 mM NaCl, 0.5% NP-40 (or its chemical equivalent IGEPAL), and 5 mg/mL Bovine Serum Albumin (BSA). Egg chambers were further blocked for 1 hour at room temperature with 10% BSA prepared in PBT. Primary antibodies diluted in “NP-40 Block” were then incubated overnight at 4°C. Samples were then washed four times with “NP-40 Block” and incubated again for at least two hours at room temperature with the secondary antibody diluted in “NP-40 Block”. After three washing steps, samples were mounted with Vectashield with DAPI (Vector Laboratories). For F-actin staining, we added Phalloidin 647 (Biolegend, 1:1000) in the second-to-last washing step. The following primary antibodies were used: mouse anti-Coracle (DSHB, 1:100). Then, the following secondary antibody was used: goat anti-mouse Alexa 568 (Invitrogen, 1:300).

### Imaging

Confocal images of fixed Drosophila BCs were collected with a 1.30 NA/63x glycerol objective on an inverted laser scanning confocal microscope Leica SP8 (Leica Microsystems, Germany) equipped with laser lines 405 (diode laser), 458, 476, 488, 496, 514 (Argon laser), 561 (diode-pumped solid-state laser), 594 and 633 nm (Helium-Neon lasers), two photomultipliers and two HyD hybrid detectors, using Las X software (Leica Mycrosystems, Germany). To score BC migration defects, images for egg chamber staging were collected with a 0.5 NA/20x dry objective on a Zeiss Axio Imager Z1 microscope (Carl Zeiss, Germany), equipped with an epi-fluorescence system, an Axiocam MR ver3.0 camera (Carl Zeiss, Germany), using Axiovision 4.9 software (Carl Zeiss, Germany). For multipoint live imaging of Drosophila BCs, egg chambers were immobilized under a “coverslip bridge” with a fibrinogen clot, as described in (Januschke and Loyer, 2020) (Chen et al., 2023). Briefly, Silicon grease was used to insulate the coverslip bridge “legs” on top of a Lumox dish (lumox® dish 35, Tissue culture dish, with foil base, Ø: 35 mm). Then, individual ovarioles without the enveloping muscle were dissected in ex vivo culture medium (Schneider’s medium (Merck) supplemented with 20% FBS (fetal bovine serum, heat inactivated; Gibco)) for no longer than 15 minutes. Ovarioles were then transferred to Schneider containing 3.5 mg/mL Fibrinogen (Sigma-Aldrich), 0.15 mg/μL insulin (Sigma-Aldrich), and 0.1 AU/μL of Thrombin (Sigma-Aldrich) to form the final clot.

Live imaging of ex vivo Drosophila BCs was performed on a Crest X-Light V3 (CrestOptics, Italy) Spinning Disk Confocal system equipped with laser lines 405, 446, 477, 518, 545, 637, and 748 nm from Lumencor (CELESTA Light Engine), a Kinetix 25 camera from Photometrics, built on an inverted Nikon ECLIPSE Ti2 microscope with a Plan Apo 60x/1.2 NA water immersion objective, using NIS elements 5.5 software (Nikon, Japan). Live imaging was performed at 25°C and BCs were imaged preferentially since delamination or the first part of migration for better optical light travel and optimal in-focus image. XY Multipoints were acquired whenever possible, time interval was variably set between 0.5-10 min and Z stacks were collected with serial optical sections separated by 0.5 to 3.5 μm. For live imaging of the actin cytoskeleton, ovarioles were stained with 3 μM SiR-actin probe (SpiroChrome, added shortly before insulin and thrombin) in Schneider-Fibrinogen for 1.5 hours before image acquisition.

### Optogenetics in BCs

For experiments with optogenetically controlled RhoGEF2 cortical recruitment, we generated flies carrying both UAS-optoGEF (UAS-pmCIBN, UAS-RhoGEF2:CRY2:mCherry (Izquierdo et al., 2018)) and UAS-RhoGAP15B::GFP and c306-Gal4 at 18°C. To prevent premature optogenetic activation, crosses and progeny were maintained in complete darkness and were handled under red light at 18°C. For in vivo optogenetic experiments, adult offspring reared at 18°C were first transferred to 29°C for 3 days to enhance Gal4 expression. Flies were then exposed to continuous blue light (427 nm LED bulb (SuperBrightLEDs)) for 6h, or kept in the darkness for 6h as controls. Ovaries were dissected immediately after treatment in the dark room under a 593 nm LED red light source (SuperBrightLEDs). After fixation and staining with Phalloidin-555 (Santa Cruz Biotechnology), BC migration ratios were evaluated as described below. For ex vivo optogenetic experiments, ovaries were dissected under a 593 nm LED red light source (SuperBrightLEDs), and activation of the light-induced pmCIBN:CRY2 interaction with concomitant membrane recruitment of RhoGEF2 was achieved using the 488 nm laser during imaging of GFP.

### Data processing and analysis

Image processing and quantification were performed with FIJI (Schindelin et al., 2012). Representative images of BC clusters are presented as either single optical section or maximum intensity projections of 2-5 planes encompassing the region of interest. For live imaging of egg chambers, image stacks were corrected for egg chamber drift using the HyperStackReg plugin (EPFL; Biomedical Imaging Group), and to improve quality of some representative images, a Denoise.ai processing step was performed within the NIS-Elements software (Nikon, Japan).

### Evaluation of RNAi-mediated depletion levels in the RhoGAP/GEF RNAi screen

Co-expression of endogenously GFP-tagged GAPs and GEFs and the UAS-driven LARIAT system (light-activated reversible inhibition by assembled trap system (Qin et al., 2017)) were used to produce high signal-to-noise fluorescent protein clusters that could be readily observed in control BC clusters but absent in cases of complete RNAi-mediated depletion. Partial RNAi depletion allowed for the detection of residual GFP puncta. In these cases, RNAi depletion was validated phenotypically (Pbl and Tum RNAi) or quantified by measuring GFP fluorescence intensity. Mean fluorescence intensity was measured in regions of interest (ROI) manually traced in the BC cluster cytoplasm (signal) as well as in a region of the egg chamber showing no specific GFP signal (background - e.g., nurse cell nuclei). Background-subtracted intensities were then averaged across five egg chambers per condition.

### Quantification of BC migration index

In fixed tissues, the extent of BC migration was quantified by calculating a migration index, defined as the ratio of elapsed to expected migration distance in Stage 10 egg chambers. Expected migration was measured by a line traced from the anterior of egg chamber to the anterior edge of the oocyte, whereas elapsed migration distance was traced from the anterior of egg chamber to the front of the BC cluster. When comparing the raw data of migration rate of controls with experimental RNAi-depleted conditions, we first performed normality tests. Given that most datasets are non-normally distributed, datasets were analysed by Mann-Whitney test and the p-values are indicated in Table S1, in Fig. 2A, B, D, and Fig. 3D. Data were plotted as stacked bars to categorize phenotypic variation.

### Quantification of BC delamination and migration velocity

Because oocyte growth is relatively consistent at a given developmental stage of the egg chamber, oocyte length at the moment of BC detachment was used as a proxy for delamination timing as in (Inaki et al., 2022). Live imaging of BC migration was used to determine the timing of cluster delamination from the follicular epithelium. Oocyte length was then measured in Fiji by tracing a line across the oocyte. For migration velocity (Fig. 3B and 3C), the position of the BC cluster was measured over time in time-lapse movies acquired every 2.5 minutes of BC clusters that had already delaminated, up to 3 hours of total imaging time, and had a maximum of 70% of the covered distance. A line was traced from the anterior pole of the egg chamber to the front of the BC cluster in each time frame, and the length was extracted. BC displacement was potted versus time, and the migration speed was calculated as the slope of the regressed line fitted in GraphPad Prism. Data distribution was tested for normality, and datasets were analyzed by the statistical test indicated in the figure legends.

For comparative analysis of migration speed upon optogenetically controlled RhoGEF2 translocation to the membrane of RhoGAP15B-overexpressing cells (Fig. 3G, 3H), we quantified the displacement from the center of the BC cluster to the oocyte. RhoGEF2 activation frequently caused a contraction of the anterior part of the egg chamber, making this an unreliable reference point. Thus, we calculated the change in the distances between the oocyte to the BC during the dark period and during the blue light period, and divided it by the elapsed time to obtain the migration speed. Analyses were restricted to BC clusters showing homogeneous UAS-RhoGAP15B-GFP expression and to clusters that had fully delaminated but were located at least 60 μm from the oocyte cortex at the start of imaging. Data distribution was tested for normality, and as the datasets passed normality tests, we used parametric paired t-tests to calculate statistical significance.

### Quantification of round BC and lead protrusion number in fixed samples

To segment the BC clusters in Figure S4, binary masks were generated from the mCD8:GFP signal using a custom FIJI macro. Inflection points along the cluster contour were detected from the binary outline, and pairs of inflection points were used as a proxy for identifying round BCs. Lead protrusion number (> 4 μm in length) forming at the migration front was quantified from the same binary images (Fig. S4B).

### Quantification of protrusion length, orientation and dynamic behaviour

For protrusion analysis we used time-lapse movies imaging the actin probe LifeAct:GFP, expressed specifically in BCs. We selected egg chambers that had migrating BC clusters. Movies were first aligned along the anterior-posterior axis. Protrusion length and orientation were then measured in 12 consecutive frames (5 min interval, 1-hour elapsed time). Protrusion orientation was determined by tracing a line from the cluster centroid to the protrusion tip. For protrusion length, we re-drawn the lines to measure the distance between the edge of the BC body to the protrusion tip. Only protrusions formed were accessed. Retracting protrusions or those incorporated in the advancing cell body were excluded. For large protrusions with multiple edges, each edge was scored independently. Protrusion dynamics were measured by tracking individual protrusions. A line was traced from the BC body to a protrusion tip in each frame.

### Quantification of myosin enrichment and spatial distribution in BC clusters

To quantify cortical myosin, BC clusters were segmented using a custom FIJI macro. Binary masks were generated from the LifeAct:GFP signal, followed by iterative erosions the define the cell perimeter. The “Analyze particles” was then used to isolate the cluster outline, and cortical regions were selected with the “Make band” tool. Myosin intensity within cortical ROIs was measured with the “measure” tool. To baseline-correct the cortical signal intensity, a portion of the nurse cell cytoplasm was used as the background signal to subtract. To generate the kymographs of myosin distribution along the front-back axis of the cluster during BC migration, we used ADAPT (Barry et al., 2015) Essentially binary masks of BC clusters were used to define the outline of the cluster (based on the “cytoplasm” and the raw Sqh:mScarlet channel was the input as the signal channel. Cortical myosin (Sqh:Scarlet) flashes were manually scored over 10 timepoints within a 12 minute timelapse sequence, and mean values are reported. To quantify myosin enrichment along the BC cortex during protrusive periods, protrusive phases were manually identified based on Lifeact:GFP signal. The corresponding timepoints in the kymographs of myosin distribution were used to generate plot profiles along the BC cluster cortex. Profiles were baseline-corrected by subtracting the minimum mean intensity value and were normalized to their respective peak intensity to reflect the relative distribution of myosin along the front-back axis of the cluster. Data were analyzed and transformed with custom FIJI macros.

### Analysis of actin flows in BCs

We used the Matlab code for PIV analysis developed in the Stramer’s team (Yolland et al., 2019). and further adapted as described in (Zhou et al., 2022). The detailed information for PIV is as follows:

1. Cell segmentation: Prior to PIV analysis, cell segmentation was performed using the Ilastik software (Berg et al., 2019). The pixel classification workflow in Ilastik was applied to generate binary masks delineating the BC clusters.
2. PIV analysis of actin flows in BCs: A 2D cross-correlation algorithm adapted from classical PIV was implemented. In brief, this method compares a region of interest in an image (source image) with a larger region of a subsequent image (search image). The sizes of the source and search regions are determined on the basis of the feature size to be tracked and the area of their expected displacement (i.e. actin bundles). For this analysis, source and search images encompassing areas of 1.4 μm2 and 2.4 μm2 were used. A cross-correlation map was computed by analyzing the cross-correlation coefficient between the source image and the search image, by shifting the source across the search one pixel at a time. Network displacement was measured by finding the maximum coefficient within the resulting cross-correlation map. To filter anomalous tracking data, only displacements that had a cross-correlation coefficient above a certain threshold, c0, were kept. For the present work, the threshold was set at c0 = 0.5. Finally, a spatial convolution with a Gaussian kernel (size of 6 μm, sigma of 1.2 μm), and temporal convolution with temporal kernel of 20 second (sigma 10 s) were used to interpolate the measured displacements to cover all the pixels within the cell outline. The complete algorithm for this analysis was implemented in Matlab.

### FRET analysis

FRET imaging was performed to monitor the spatiotemporal dynamics of Rho activity in migrating BC clusters. Flies expressing the Raichu-RhoA FRET biosensor ((Itoh et al., 2002; Qin et al., 2017; Yoshizaki et al., 2003)) were dissected and mounted for live imaging as previously described. Images were acquired using a Zeiss LSM 710 confocal microscope equipped with a 63×/1.40 NA oil-immersion objective and controlled by ZEN software.

The donor fluorophore (CFP) was excited with a 440-nm laser line, and emission signals were collected sequentially in the donor channel (460–500 nm) and FRET channel (520–560 nm) using separate detection tracks to minimize cross-excitation. Reference images of samples expressing donor-only or acceptor-only constructs were obtained under identical settings to calculate bleed-through correction coefficients.

Corrected FRET efficiency was expressed as the FRET/CFP emission ratio, and pseudo colored ratio images representing relative Rho activity were generated using ImageJ with custom-written macros. Ratio values were normalized to the mean activity of follower BCs within each cluster to obtain the relative Rho activity values shown in figures.

Time-lapse FRET imaging was conducted at 1 min intervals for 15–20 min at room temperature. Identical acquisition parameters were used across all genotypes, including control and RhoGAP15B RNAi clusters. Data was obtained from at least three independent experiments. Statistical analyses were performed using GraphPad Prism, and differences between groups were evaluated using an unpaired Student’s *t*-test. Increased FRET ratios indicate higher Rho activation. FRET imaging was performed under identical settings for control and RhoGAP15B RNAi BC groups.

### Statistical data analysis

Statistical analysis and graphs were generated using GraphPad Prism 9 (GraphPad Software, La Jolla, CA, USA). All fly crosses were repeated at least twice. Sample sizes indicated (n) in figure legends refer to egg chambers. For analysis of fixed tissues, from 6 to 10 flies were dissected per independent experiment, whereas 2 to 3 BC clusters were imaged per dissected ovary for live imaging of BC migration. If data were not normally distributed and two groups were compared, an unpaired, Mann-Whitney U test was used for statistical analysis. If data were normally distributed and two groups were compared, a student’s t-test was used for statistical analysis.

## Author Contributions

E.M. and V.Y. conceptualized the study and wrote the original manuscript draft. Data acquisition: V.Y., C.M., M.O. and E.M. performed and analyzed most experiments with exception of FRET and PIV analysis, which were performed by B.L. and X.W. J.L. and Y.B generated RhoGAP15B mutant lines. Visualization: E.M and V.Y. Data analysis and interpretation: V.Y., C.M., B.L., X.W and E.M. Supervision: Y.B., X.W. and E.M. Funding acquisition: Y.B., X.W and E.M. All authors reviewed the manuscript.

## Acknowledgments

The authors thank the Bloomington Drosophila Stock Center (BDSC) and the Vienna Drosophila Resource Center (VDRC) for fly stocks. Work in the E.M. lab is funded by National Funds through FCT—Fundação para a Ciência e a Tecnologia, I.P., under the projects PTDC/BIA-CEL/1511/2021 and UIDB/04293/2020. V. Yang was supported by a PhD Fellowship (2020.05791.BD) from the FCT - Fundação para a Ciência e Tecnologia, IP. The authors also acknowledge the support of the i3S Scientific Platform ALM, a member of the national infrastructure PPBI-Portuguese Platform of Bioimaging (PPBI-POCI-01-0145-FEDER-022122). Work in the X. Wang lab is supported by Agence Nationale de la Recherche (ANR PRC AAPG2022 NEW-corset) and Scientifiques de la Fondation ARC (PJA20191209714)

Work in the Y.B. lab is supported by the Institut Curie, the CNRS, the INSERM as well as ARC (SL220130607097), ANR (TiMecaDiv 20CE13000801), CANCERO-INCA (PLBIO2020/BELLAICHE) grants.

## Notes

### Competing Interest Statement

The authors have declared no competing interest.

