## Supplementary material for "Spatial inhibition of RhoA by RhoGAP15B promotes protrusive activity during collective migration": Table S2

**Table S2. Key Antibodies and Chemicals**

| Name/Description | Source | Number |
|---|---|---|
| Tween 20 | Merck | P9416 |
| Paraformaldehyde | em-grade.com | GF750170 |
| Phalloidin 555 | Santa Cruz Biotechnology | sc-363794 |
| Phalloidin 647 | Biolegend | 424205 |
| Vectashield with DAPI | Vector Laboratories | H-1200 |
| Schneider's Insect Medium | Merck | S0146 |
| Fetal Bovine Serum | Gibco | A5256801 |
| Fibrinogen | Sigma-Aldrich | F8630 |
| Insulin | Sigma-Aldrich | I0516 |
| Thrombin | Sigma-Aldrich | T7513 |
| SiR-actin | SpiroChrome | SC001 |
| Coracle antibody (mouse) | Developmental Studies Hybridoma Bank (DSHB) | RRID:AB_1161644 |
| Goat anti-mouse Alexa 568 | Invitrogen | RRID:AB_144696 |
| IGEPAL | Sigma-Aldrich | I3021 |
| Tris base | Calbiochem | 648311 |
| Hydrochloric acid (HCl) 37% | Emsure/Merck | 1.00317 |
| Sodium Chloride (NaCl) | NZYTech | MB15901 |
| Bovine Serum Albumin (BSA) | Sigma-Aldrich | A7906 |
