## Supplementary material for "Spatial inhibition of RhoA by RhoGAP15B promotes protrusive activity during collective migration": Table S3

**Table S3. *Drosophila melanogaster* Stocks.**
BDSC - Bloomington Drosophila Stock Center; KDSC - Kyoto Drosophila Stock Center; VDRC - Vienna Drosophila Resource Center.

| **Stock description and genotype** | **Source** | **Identifier** |
|---|---|---|
| Tumbleweed:GFP | Di Pietro et al. 2023 | N/A |
| Pebble:GFP | Di Pietro et al. 2023 | N/A |
| RhoGAP15B:GFP | Di Pietro et al. 2023 | N/A |
| Sos:GFP | Di Pietro et al. 2023 | N/A |
| Zir:GFP | Di Pietro et al. 2023 | N/A |
| Trio:GFP | Di Pietro et al. 2023 | N/A |
| Graf:GFP | Di Pietro et al. 2023 | N/A |
| RhoGAP19D:GFP | Di Pietro et al. 2023 | N/A |
| cdGAPr:GFP | Di Pietro et al. 2023 | N/A |
| CG43658:GFP | Di Pietro et al. 2023 | N/A |
| Conu:GFP | Di Pietro et al. 2023 | N/A |
| RtGEF:GFP | Di Pietro et al. 2023 | N/A |

| | | |
|---|---|---|
| RhoGAP1A:GFP | Di Pietro et al. 2023 | N/A |
| RhoGAPp190:GFP | Di Pietro et al. 2023 | N/A |
| RhoGAP68F:GFP | Di Pietro et al. 2023 | N/A |
| Cysts:GFP | Di Pietro et al. 2023 | N/A |
| Ecad:GFP | Huang et al., 2009 | N/A |
| aPKC:GFP | Chen et al., 2018 | N/A |
| slbo>LifeAct:GFP | BDSC | RRID: VDRC_ 58363 |
| *Gr1*-Gal4 | BDSC | RRID: BDSC_36287 |
| *tj*-Gal4 | KDSC | RRID: KDSC_104055 |
| *c306*-Gal4 | BDSC | RRID: BDSC_3743 |
| *slbo*-GAL4 | BDSC | RRID: BDSC_ 58435 |
| Gal80$^{ts}$ | BDSC | RRID: BDSC_7017 |

| | | |
|---|---|---|
| UAS-*trio* RNAi | BDSC | RRID: BDSC_27732 |
| UAS-*cysts* RNAi | BDSC | RRID: BDSC_ 33047 |
| UAS-*CG43658* RNAi | BDSC | RRID: BDSC_ 32341 |
| UAS-*Zir* RNAi | BDSC | RRID: BDSC_ 28005 |
| UAS-*Sos* RNAi | BDSC | RRID: BDSC_ 34833 |
| UAS-*RtGEF* RNAi | VDRC | RRID: VDRC_ 105093 |
| UAS-*Vav* RNAi | BDSC | RRID: BDSC_ 39059 |
| UAS-*Pbl* RNAi | BDSC | RRID: BDSC_ 28343 |
| UAS-*Conu* RNAi | VDRC | RRID: VDRC_ 104232 |
| UAS-*Graf* RNAi | BDSC | RRID: BDSC_ 51853 |
| UAS-*RhoGAP15B* RNAi | BDSC | RRID: BDSC_ 6427 |
| UAS-*RhoGAP15B* RNAi | BDSC | RRID: BDSC_ 42527 |

| | | |
|---|---|---|
| UAS-*RhoGAP15B* RNAi | VDRC | RRID: VDRC_ 24678 |
| UAS-*RhoGAPp190* RNAi | BDSC | RRID: BDSC_ 43987 |
| UAS-*RhoGAP68F* RNAi | BDSC | RRID: BDSC_ 6442 |
| UAS-*Tum* RNAi | BDSC | RRID: BDSC_ 28928 |
| UAS-*RhoGAP19D* RNAi | BDSC | RRID: BDSC_ 6436 |
| UAS-*RhoGAPp1A* RNAi | BDSC | RRID: BDSC_ 33390 |
| UAS-*CdGAPr* RNAi | BDSC | RRID: BDSC_ 38279 |
| UAS-*mCherry* RNAi | BDSC | RRID: BDSC_ 35785 |
| $w^{1118}$ | BDSC | RRID: BDSC_3605 |
| $rhoGAP15B^{237}$ | This paper | N/A |
| FRT19A $rhoGAP15B^{237}$/ nls:RFP hsFLP FRT19A (X) | This paper | N/A |
| UAS-LARIAT | Qin et al., 2017 | N/A |

| | | |
|---|---|---|
| UAS-RhoGAP15B:HA | This paper | N/A |
| UAS-RhoGAP15B:GFP | This paper | N/A |
| UAS-CIBN-CAAX | Izquierdo et al., 2018 | N/A |
| UAS-RhoGEF2:CRY2:mCherry | Izquierdo et al., 2018 | N/A |
| UAS-Utrophin:RFP | Herszterg et al., 2013 | N/A |
| UAS-RhoAFRET | Qin et al., 2017 | N/A |
| UAS-mCD8:GFP | BDSC | RRID: BDSC_ 76363 |
| UAS-Rho1.V14 (CA) | BDSC | RRID: BDSC_ 7330 |
