## Supplementary material for "Spatial inhibition of RhoA by RhoGAP15B promotes protrusive activity during collective migration": Table S4

**Table S4. List of Fly Genotypes and experimental conditions**

| **Figure** | **Condition** |
| --- | --- |
| 1 | endogenously RhoGAP/GEF GFP-tagged lines (See Table S2) |
| 2A | *tj-Gal4*/UAS-LARIAT; Trio:GFP/UAS-*mCh*RNAi,  3 days 29 °C  *tj-Gal4*/UAS-LARIAT; Trio:GFP/UAS-*Trio*RNAi,  3 days 29 °C  Cysts:GFP/+; *GR1-Gal4,* UAS-LARIAT/+,  3 days 29 °C  Cysts:GFP/+; *GR1-Gal4,* UAS-LARIAT/UAS-*Cysts*RNAi,  3 days 29 °C  *tj-Gal4*/UAS-LARIAT; CG43658:GFP/UAS-*mCh*RNAi,  3 days 29 °C  *tj-Gal4*/UAS-LARIAT; CG43658:GFP/UAS-*CG43658*RNAi,  3 days 29 °C  Zir:GFP/+; *GR1-Gal4,* UAS-LARIAT/+,  3 days 29 °C  Zir:GFP/+; GR1-Gal4, UAS-LARIAT/UAS-*Zir*RNAi,  3 days 29 °C  Sos:GFP/+; *GR1-Gal4,* UAS-LARIAT/+,  7 days 25 °C  Sos:GFP/+; *GR1-Gal4*, UAS-LARIAT/UAS-*Sos*RNAi,  7 days 25 °C  RtGEF:GFP/ *GR1-Gal4*, UAS-LARIAT,  3 days 29 °C  UAS-*RtGEF*RNAi/+; *GR1-Gal4*,UAS-LARIAT/RtGEF:GFP,  3 days 29 °C  Vav:GFP/+;; *GR1-Gal4*,UAS-LARIAT/UAS-*Vav*RNAi,  3 days 29 °C  Vav:GFP/+;; *GR1-Gal4*,UAS-LARIAT/+,  3 days 29 °C  tj-Gal4/CyO; Pebble:GFP/UAS-LARIAT,  7 days 25 °C  tj-Gal4/UAS-LARIAT; Pebble:GFP/UAS-*Pebble*RNAi,  7 days 25 °C |
| 2B | Conu:GFP/+; *GR1-Gal4*, UAS-LARIAT/UAS-*mCh*RNAi,  7 days 25 °CG  Conu:GFP/+; *GR1-Gal4*, UAS-LARIAT/UAS-*Conu*RNAi,  7 days 25 °C  Graf:GFP/+; UAS-*Graf*RNAi/+; *GR1-Gal4*, UAS-LARIAT/+,  3 days 29 °C  Graf:GFP/+; +/+; *GR1-Gal4*, UAS-LARIAT/+,  3 days 29 °C  RhoGAP15B:GFP/+; +/+; *GR1-Gal4*, UAS-LARIAT/+,  3 days 29 °C  RhoGAP15B:GFP/+; +/+; *GR1-Gal4*, UAS-LARIAT/UAS-*RhoGAP15B*RNAi (BDSC_6427)  3 days 29 °C  RhoGAPp190:GFP/+; +/+; *GR1-Gal4*, UAS-LARIAT/+,  3 days 29 °C  RhoGAPp190:GFP/+; UAS-*RhoGAPp190*RNAi/+; *GR1-Gal4*,UAS-LARIAT/+,  3 days 29 °C  *GR1-Gal4*, UAS-LARIAT/RhoGAP68F:GFP,  3 days 29 °C  UAS-*RhoGAP68F*RNAi/+; *GR1-Gal4*, UAS-LARIAT/RhoGAP68F:GFP,  3 days 29 °C  Tum:GFP/+; *GR1-Gal4*, UAS-LARIAT/+,  3 days 29 °C  Tum:GFP/+; *GR1-Gal4*, UAS-LARIAT/UAS-*Tum*RNAi,  3 days 29 °C  RhoGAP19D:GFP/+;; *GR1-Gal4*, UAS-LARIAT/+,  7 days 25 °C  RhoGAP19D:GFP/+;; *GR1-Gal4,* UAS-LARIAT/UAS-*RhoGAP19D*RNAi,  7 days 25 °C  RhoGAP1A:GFP/+;; *GR1-Gal4*, UAS-LARIAT/+,  3 days 29 °C  RhoGAP1A:GFP/+;; *GR1-Gal4*, UAS-LARIAT/UAS-*RhoGAP1A*RNAi,  3 days 29 °C  CdGAPr:GFP/+; *GR1-Gal4*, UAS-LARIAT/UAS-*mCh*RNAi,  7 days 25 °C  CdGAPr:GFP/UAS-*CdGAPr*RNAi; *GR1-Gal4*,UAS-LARIAT/+  7 days 25 °C |
| 2C | RhoGAP15B:GFP/+; ; *GR1-Gal4*,UAS-LARIAT/+,  3 days 29 °C  RhoGAP15B:GFP/+; ; *GR1-Gal4*,UAS-LARIAT/UAS-*RhoGAP15B*RNAi,  3 days 29 °C |
| 2D | c306-Gal4/+; +/+; Gal80^ts^/UAS-*mCh*RNAi,  c306-Gal4/+; UAS-*RhoGAP15B*RNAi(VDRC_24678)/+; Gal80^ts^/+,  c306-Gal4/+; UAS-*RhoGAP15B*RNAi(BDSC_42527)/+; Gal80^ts^/+,  *upd-Gal4*/+; +/+; UAS-*mCh*RNAi/+,  *upd-Gal4*/+; UAS-*RhoGAP15B*RNAi(VDRC_24678),  FRT19A,  FRT19A *rhoGAP15B^237^* |
| 2E | FRT19A  FRT19A *rhoGAP15B^237^* |
| 3A-3C | c306-Gal4/+;; slbo>LifeAct:GFP/UAS-*mCh*RNAi,  c306-Gal4/+; UAS-*RhoGAP15B*RNAi(VDRC_24678)/+; slbo>LifeAct:GFP/+,  FRT19A, Δ*rhoGAP15B^237^*;; slbo>LifeAct:GFP /+ |
| 3D-3H | c306-Gal4/+;; UAS-mCherry-RNAi/+,  c306-Gal4/+;; UAS-RhoGAP15B:GFP/+,  c306-Gal4/UAS-CIBN-CAAX;; UAS-RhoGAP15B:GFP/UAS-mCherry-RNAi,  c306-Gal4/UAS-CIBN-CAAX;; UAS-RhoGAP15B:GFP/UAS- RhoGEF:CRY2:mCherry |
| 4A | RhoGAP15B:GFP |
| 4B | RhoGAP15B:GFP;; slbo-Gal4,UAS-Utrophin:RFP |
| 4C-4G | c306-Gal4/+;; slbo >LifeAct:GFP/UAS-*mCh*RNAi,  c306-Gal4/+; UAS-*RhoGAP15B*RNAi(VDRC_24678)/+; slbo-UAS-LifeAct:GFP/+  c306-Gal4/+;; slbo >LifeAct:GFP/UAS-RhoGAP15B-HA |
| 5A-5C | *c306-Gal4*/+ ; UAS-RhoFRET/+ ; UAS-RhoFRET/UAS-*mCh*RNAi,  c306-Gal4/+ ; UAS-RhoFRET/UAS-*RhoGAP15B*RNAi(VDRC_24678) ; UAS-RhoFRET/+ |
| 5D-5J | c306-Gal4/+; +/+; slbo-LifeAct:GFP/UAS-*mCh*RNAi,  c306-Gal4/+; UAS-*RhoGAP15B*RNAi(VDRC_24678)/+; slbo>LifeAct:GFP/+,  c306-Gal4/+;; UAS-RhoGAP15B-HA/+ |
| S1 | endogenously RhoGAP/GEF GFP-tagged lines (di Pietro et al., 2023) |
| S2B | Cysts:GFP/+; *GR1-Gal4,* UAS-LARIAT/+,  Cysts:GFP/+; *GR1-Gal4,* UAS-LARIAT/UAS-*Cysts*RNAi |
| S2C | *tj-Gal4*/UAS-LARIAT; CG43658:GFP/UAS-*mCh*RNAi,  *tj-Gal4*/UAS-LARIAT; CG43658:GFP/UAS-*CG43658*RNAi |
| S2D | Zir:GFP/+; *GR1-Gal4,* UAS-LARIAT/+,  Zir:GFP/+; GR1-Gal4, UAS-LARIAT/UAS-*Zir*RNAi |
| S2E | *tj-Gal4*/UAS-LARIAT; CG43658:GFP/UAS-*mCh*RNAi,  *tj-Gal4*/UAS-LARIAT; CG43658:GFP/UAS-*CG43658*RNAi |
| S2F | *tj-Gal4*/CyO; Pebble:GFP/UAS-LARIAT,  *tj-Gal4*/UAS-LARIAT; Pebble:GFP/UAS-*Pebble*RNAi |
| S2G | Conu:GFP/+; *GR1-Gal4*, UAS-LARIAT/UAS-*mCh*RNAi,  Conu:GFP/+; *GR1-Gal4*, UAS-LARIAT/UAS-*Conu*RNAi |
| S2H | Graf:GFP/+; UAS-*Graf*RNAi/+; *GR1-Gal4*,UAS-LARIAT/+,  Graf:GFP/+; +/+; *GR1-Gal4*,UAS-LARIAT/+ |
| S2I | Tum:GFP/+; *GR1-Gal4*,UAS-LARIAT/+,  Tum:GFP/+; *GR1-Gal4*,UAS-LARIAT/UAS-*Tum*RNAi |
| S2J | CdGAPr:GFP/+; *GR1-Gal4*,UAS-LARIAT/UAS-*mCh*RNAi,  CdGAPr:GFP/UAS-*CdGAPr*RNAi; *GR1-Gal4*,UAS-LARIAT/+ |
| S3A | RhoGAP15B:GFP |
| S3B-S3D | FRT19A, *rhoGAP15B^237^*/nls:RFP,hsFLP,FRT19A; E-Cad:GFP/+ |
| S3E | FRT19A, *rhoGAP15B^237^*/nls:RFP,hsFLP,FRT19A; aPKC:GFP/+ |
| S4 | +/+; slbo-Gal4,UAS-mCD8:GFP/+,  FRT19A,Δ*rhoGAP15B^237^*; slbo-Gal4,UAS-mCD8:GFP/+,  slbo-Gal4,UAS-mCD8:GFP/UAS-Rho.CA |
