## Supplemental Figures and Movie Legends for "Spatial inhibition of RhoA by RhoGAP15B promotes protrusive activity during collective migration"

Supplementary DATA file includes 4 Figures and Movie Legends

**Figure S1. Localization of RhoGAPs and RhoGEFs during delamination and neodelamination.**

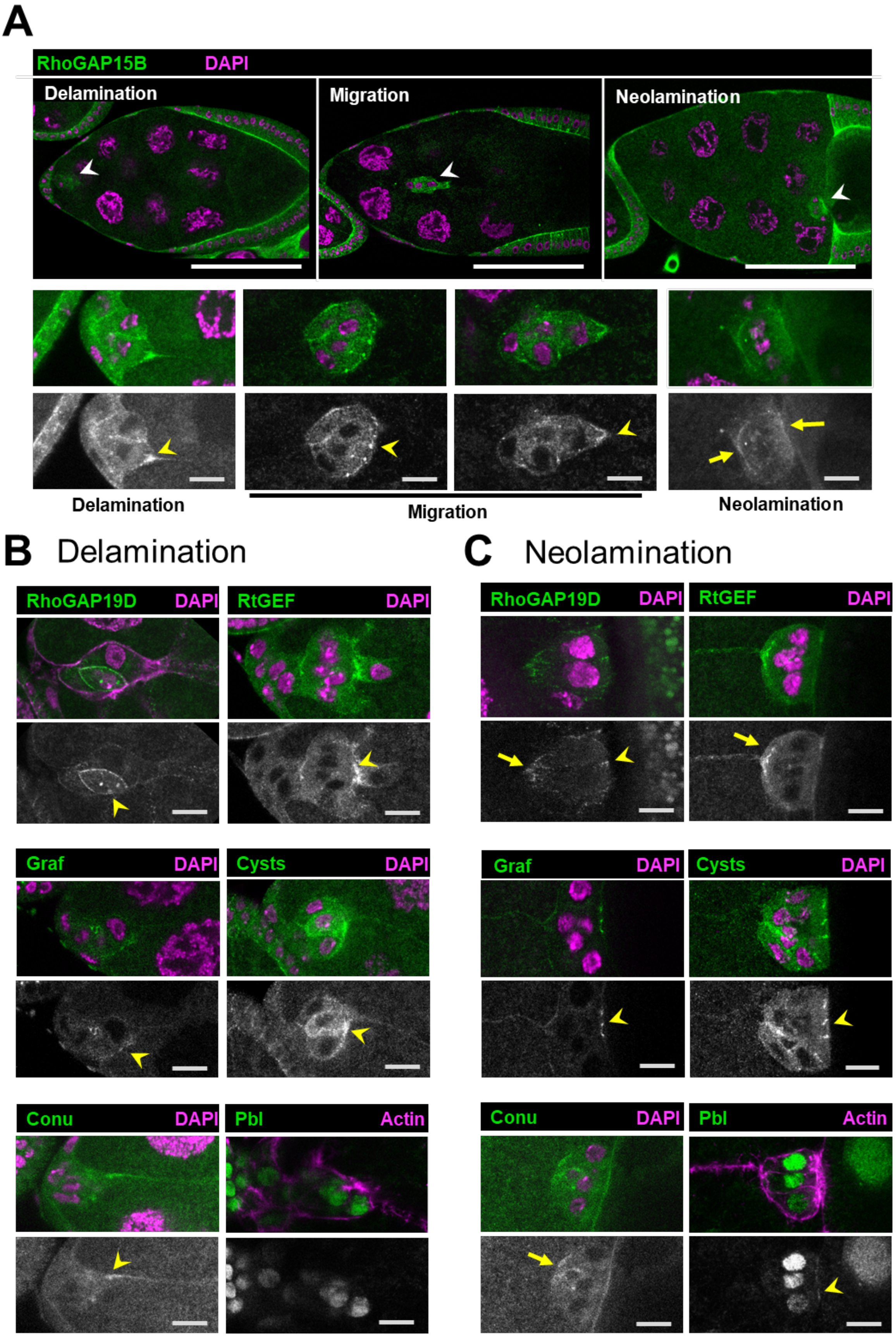

(**A)** Confocal images of RhoGAP15B:GFP egg chambers illustrating the three stages of BC migration. RhoGAP15B:GFP shows local enrichment at the migration front (yellow arrowheads) since delamination (left, close-ups). During migration is observed in puncta along the outer cortex, and displays front accumulation prior to protrusion formation (center, close-ups). During the neolamination phase, enrichment appears at both the soon-to-be basal (left yellow arrow) and apical (right yellow arrow), consistent with the localization in follicle cells.
**(B and C)** Confocal images of endogenously GFP-tagged RhoGAPs and RhoGEFs showing relevant subcellular localization patterns during delamination (B) and neolamination (C). (B) During delamination, RhoGAP19D shows strong accumulation at the polar-BC contact (yellow arrowheads), RtGEF accumulates at the rear cortex of the leading cell (yellow arrowheads), and Cysts and Conu display local enrichment at the BC cortex (yellow arrowheads). (C) During neolamination, RhoGAP19D, Cysts and Graf accumulate at intercellular junctions near the apical side (yellow arrowheads), whereas Pbl shows broader apical enrichment (yellow arrowheads). RhoGAP19D, Conu and RtGEF also display strong enrichment at the rear of the cluster (yellow arrows in (C) (A-C) Egg chambers were stained with DAPI (nuclei) or phalloidin (Actin) as indicated (magenta). (A-C) GFP signals are also shown separately (bottom panels). Scale bars: 100 µm (wide view of egg chambers (A)) and 10 µm (BC close-ups (A-C)).

**Figure S2. RhoGAP/RhoGEFs showing delayed BC migration defects**

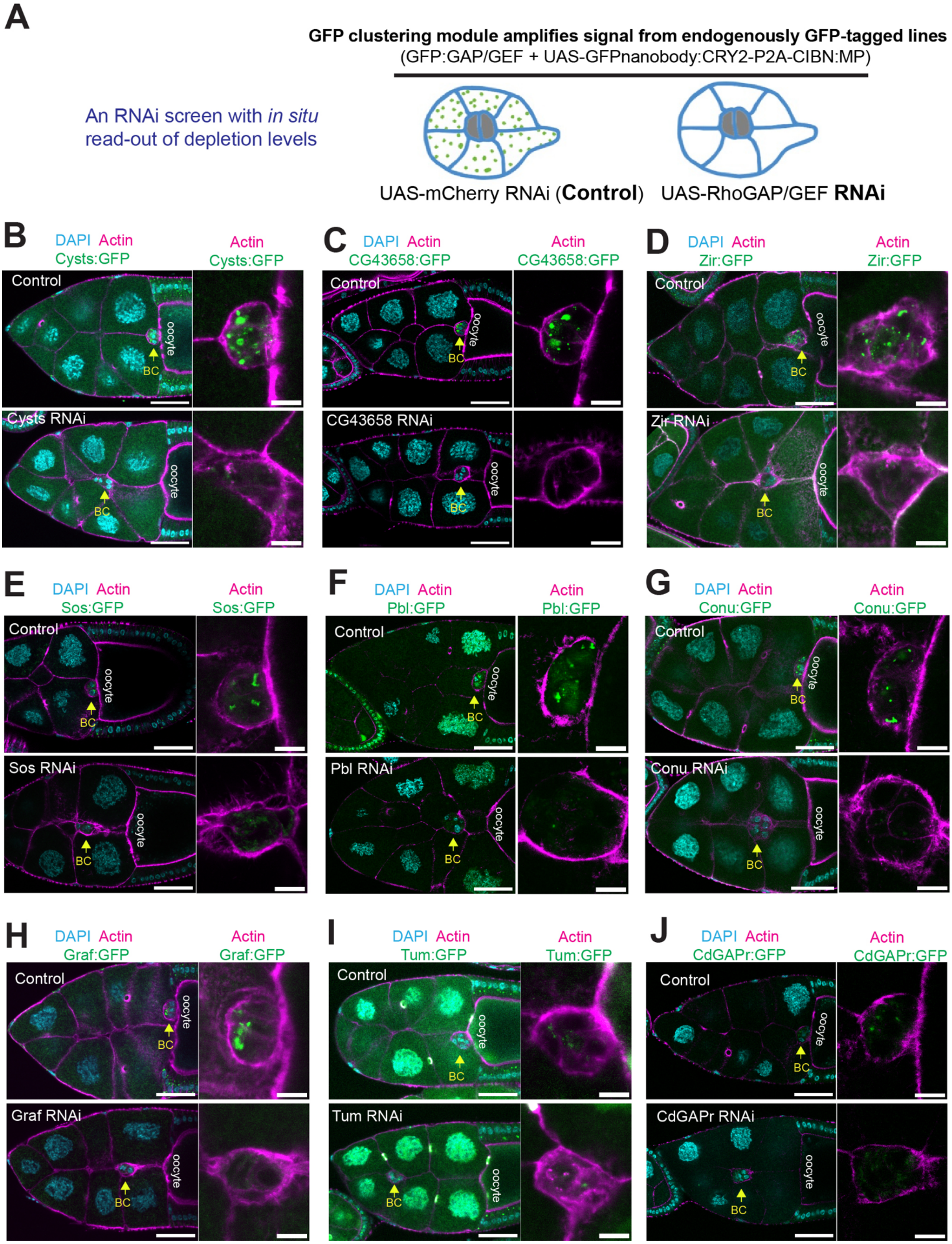

**(A)** Schematic of the RNAi screen coupled with depletion validation. UAS-driven RNAi lines and respective controls were co-expressed with the corresponding endogenously GFP-tagged protein and a signal amplification module based on light-induced GFP clustering (UAS-GFPnanobody:CRY2-P2A-CIBN:MP (Qin et al., 2017)).
**(B–J)** Representative images of delayed BC migration after depletion of Cysts (**B**), CG43658 (**C**), Zir (**D**), Sos (**E**), Pebble (**F**), Conu (**G**), Graf (**H**), Tum (**I**), and CdGAPr (**J**) (green). Yellow arrows indicate BC clusters. Egg chambers co-expressing the corresponding endogenously GFP-tagged line (green) and the GFP clustering module were stained for F-actin (magenta) and the nuclei (DAPI, cyan). Enlarged views of the BC clusters confirm depletion. Arrows indicate BC position. Scale bars: wide view - 50 µm; close-ups - 10 µm.

**Figure S3. RhoGAP15B is not required for BC apical-basal polarity and adhesion**

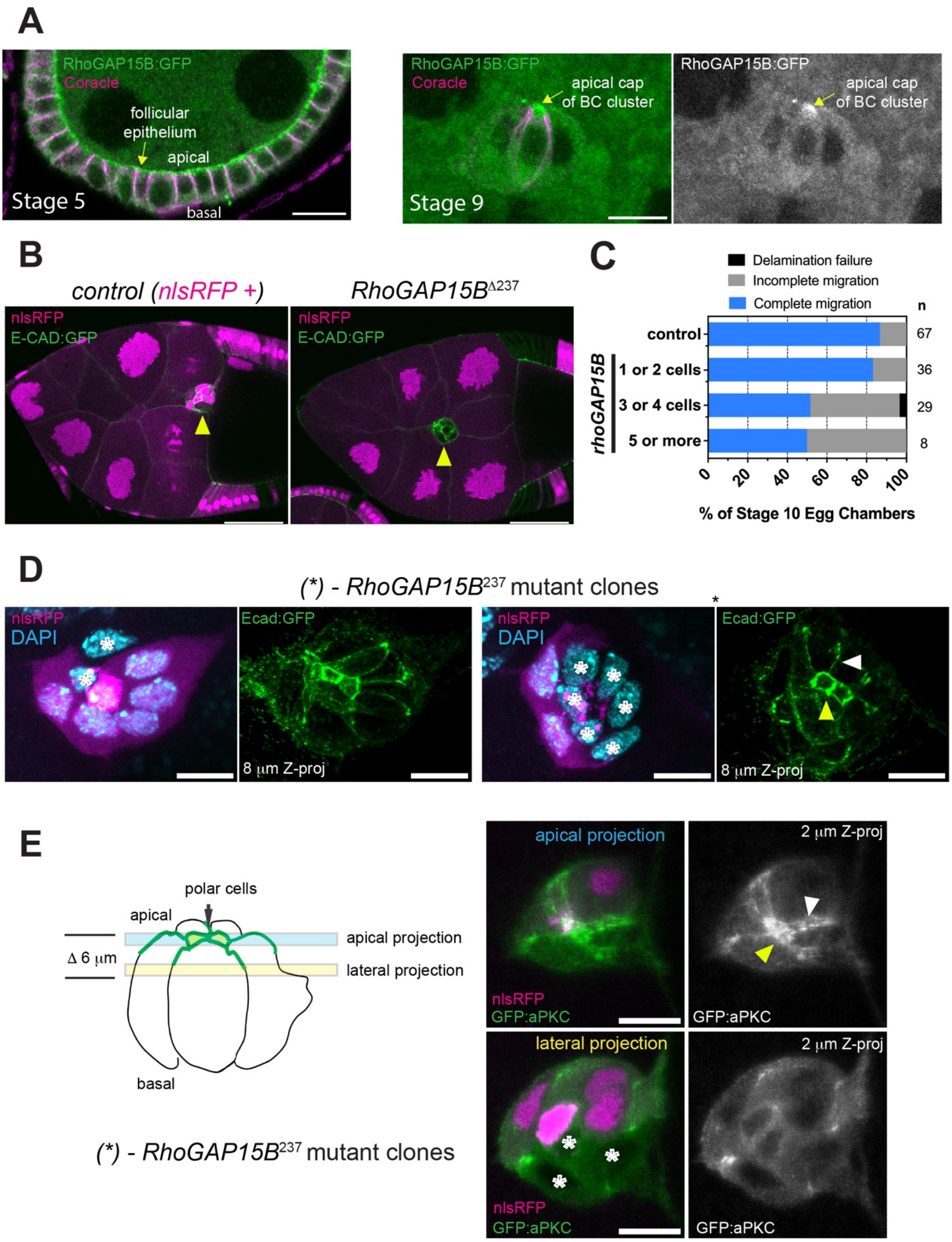

**(A)** Immunofluorescence of Coracle (magenta) co-imaged with endogenously tagged RhoGAP15B:GFP (green). RhoGAP15B accumulates apically in the follicular epithelium (left) and at the apical cap of polar cells within the BC cluster (right). Coracle labels the lateral cortex and the polar cell septate junctions. Scale bar, 10 µm.
**(B and C)** BC clusters containing clonal *RhoGAP15B* mutant cells in a wild-type germline show delayed migration (B). Quantification indicates that migration defects are only frequent when at least half the cluster is mutant for *RhoGAP15B (C)*. nlsRFP (magenta) marks wild-type clonal cells. *n* indicates the number of stage 10 egg chambers analyzed. Scale bar, 50 µm.
**(D)** Top view maximum-intensity projections (8 µm) of BC clusters containing clonal *RhoGAP15B* mutant cells show no effect on E-cadherin localization (green) at BC junctions (white arrowhead) or polar cell junctions (yellow arrowhead). Asterisks indicate mutant cells. Nuclei were stained with DAPI (cyan). Scale bar, 10 µm.
**(E)** Top view maximum-intensity projections of the apical region (top) and lateral (bottom) regions of BC clusters containing clonal *RhoGAP15B* mutant cells show that *RhoGAP15B* mutation does not affect aPKC localization at the apical cap (yellow arrow) or at the apical BC-BC contact in comparison to control cells (marked with nlsRFP). Scale bar, 10 µm.

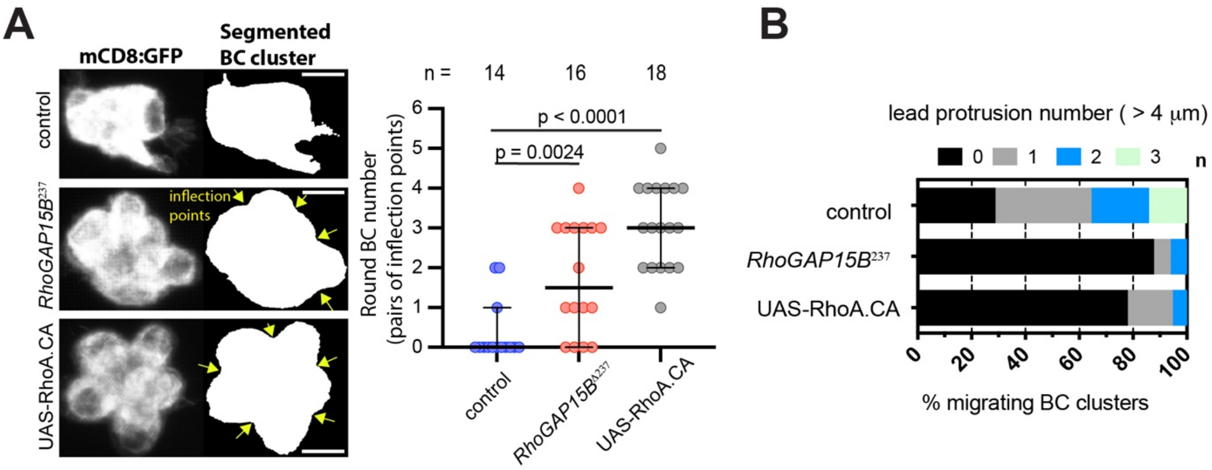


**Figure S4. *RhoGAP15B* mutants mimic RhoA activation phenotypes in BC morphology**

**(A)** Morphometric analysis of BC clusters in controls, *RhoGAP15B* mutants, and Rho.CA OE co-expressing UAS-mCD8:GFP with slbo-Gal4 to segment BCs. Yellow arrows indicate inflection points (concave regions flanked by convex surfaces), used to quantify rounded BC number. Plot shows quantification of rounded cell number. Data are presented as median +/- 95% confidence interval. Statistical significance was assessed using one-way ANOVA. **(B)** Frequency distribution of lead protrusion number detected within the migration front (-45º to 45º) in controls, *RhoGAP15B* mutants, and Rho.CA OE fixed egg chambers. Scale bars: 10 µm.

### Supplementary movie legends

**Movie S1) Delayed BC migration by genetic interference with RhoGAP15B**

Time-lapse video of a stage 9 egg chamber expressing slbo>LifeAct:GFP (BCs, green) in control (top), c306>Gal4 driven RhoGAP15B RNAi (middle) and *RhoGAP15B*$^{239}$ mutant (bottom). Time interval: 5 min. Scale bar: 20 μm.

**Movie S2) Light-induced cortical recruitment of a RhoGEF restores migration speed of RhoGAP15B-overexpressing BCs**

Time-lapse video of a stage 9 egg chamber expressing c306-Gal4>UAS-RhoGAP15B:GFP (OE, green) with (bottom) or without (top) co-expression of UAS-CAAX:CIBN and UAS-CRY2:RhoGEF2, and stained with the live actin probe SiR-Act (magenta). Imaging with 488 nm laser triggers RhoGEF2 cortical recruitment from time 0 onward. Time interval: 0.5 min. Scale bar: 20 μm.

**Movie S3) 3D rendering comparing the morphology of BC clusters mutant for *RhoGAP15B* or overexpressing constitutively active RhoA**

Fixed stage 9 BCs expressing *slbo*-Gal4>UAS-mCD8:GFP (3D projection, green) and a z-slice segmentation of BCs (3D reconstruction, gray); Video shows control (left), *RhoGAP15B*$^{239}$ mutant (middle) and UAS-RhoA-CA (right). Scale bar: 10 μm.

**Movie S4) RhoGAP15B regulates front protrusions dynamics**

Time-lapse video of slbo>LifeAct:GFP (inverted gray LUT) stage 9 control (top) egg chambers, and egg chambers depleted of *RhoGAP15B* (RNAi middle) or overexpressing UAS-RhoGAP15B (OE, bottom). The time interval is 2.5 min. Scale bar: 20 μm.

**Movie S5) Migrating *RhoGAP15B* RNAi BC clusters show frequent myosin pulses at the front**

Time-lapse video of stage 9 egg chamber expressing squash:mScarlet (fire LUT) and slbo>LifeAct:GFP (gray). Left is control and right is RhoGAP15B RNAi. The time interval is 10 sec. Scale bar: 10 μm.
